# A Creb3-Like Transcription Factor Coordinates ER Function upon Food Intake to Regulate Lipid Metabolism

**DOI:** 10.1101/2021.03.13.435227

**Authors:** Haris A. Khan, Ming Toh, Tamás Schauer, Rory J. Beresford, Paula Ortega-Prieto, Catherine Postic, Carla E. Margulies

## Abstract

Ingestion of nutrients elicits essential physiological responses, including absorption, digestion, cessation of feeding and nutrient storage. The endoplasmic reticulum (ER) is central to this nutritional homeostasis, since it regulates intracellular organelle function, drives intercellular communication and promotes metabolite distribution. We identified the *Drosophila* Creb3L-family transcription factor, CrebA, as the key metabolic regulator of ER function, thereby affecting lipid metabolism and feeding behavior. In response to feeding, CrebA activity is rapidly and transiently activated. CrebA directly drives the expression of the ER protein sorting machinery. We demonstrate that CrebA levels regulate lipid metabolism through lipoprotein secretion into the hemolymph and suppress feeding behavior. Further, CrebA mouse homologs are also upregulated in the liver following feeding and drive the transcriptional activation of ER protein sorting machinery genes in mammals. Our results reveal an evolutionarily conserved transcription switch which is turned on in response to food ingestion and orchestrates a negative feedback loop that promotes satiety by regulating ER function and protein secretion.

## INTRODUCTION

Regulating nutrient intake is a basic process of living organisms, helping to ensure that individuals have the appropriate building blocks and energy to function in ever-changing environments. The ER plays a pivotal role in the communication and metabolite redistribution necessary for survival. Cells in higher organisms report their nutritional status by secreting hormones and sensing these hormones *via* cellular membrane receptors. Further, organisms redistribute metabolites between cells, either by regulating their secretion or uptake. ER function is thus crucial in the cells of metabolically highly active organs such as the liver.

The ER also organizes intracellular metabolism. ER morphology and function depend on nutritional state (Cruz-Garcia et al., 2018; van Leeuwen and Rabouille, 2019) and are affected in metabolic diseases (Huang et al., 2019; Nishi et al., 2019). In addition to signaling peptides, which are synthesized and processed in the ER and Golgi apparatus, proteins required for sugar and lipid metabolism also depend on ER function. The production of glucose from glucose-6-phosphate in the liver, for example, occurs in the ER (Waddell and Burchell, 1991; Hutton and O’Brien, 2009), as does the synthesis of triacylglycerols (Coleman et al., 2000), phospholipids (Fagone and Jackowski, 2009) and cholesterol esters (Chang et al., 2006). Since the metabolic decision to form cellular lipid droplets or secrete vascular lipoprotein particles is made in the ER (Yao et al., 2013; Gillon et al., 2012; Koerner et al., 2019; Balla et al., 2019), its regulation is key to metabolic health (Baiceanu et al., 2016). Although we have a broad understanding of lipid metabolism and protein secretion, the cellular decisions that link nutrient uptake with ER function and lipid homeostasis is not well understood at the molecular level.

Mobilizing lipids in aqueous milieus is difficult. Thus, metazoans evolved mechanisms to shuttle lipids between cells in the form of lipoprotein particles. Both vertebrates and insects depend on ER function for lipid transport (Sirwi and Hussain, 2018). In vertebrates, the major component of these particles includes ApoA, which provides the protein scaffold for high-density lipoproteins (HDL), and ApoB, which provides the scaffold for chylomicrons and very low-density lipoproteins (VLDL). Flies have two major proteins, Apolipophorin (Apolpp) and Lipid transfer particle (Apoltp), which show homology to the N-terminal domain of vertebrate ApoB. ApoB is synthesized into the lumen of the ER, where the Microsomal triglyceride transfer protein (MTP) transfers lipids to ApoB (Hussain et al., 2003; Sirwi and Hussain, 2018). Absence of MTP in humans and flies results in the absence of circulating ApoB (Aggerbeck et al., 1992) and Apolpp/Apoltp, respectively (Palm et al., 2012). For further maturation, these particles exit the ER and are transported to the Golgi *via* specialized COPII vesicles which carry large cargos (Santos et al., 2016; Siddiqi et al., 2018; Melville et al., 2020). Importantly, interfering with this process prevents lipid transport from the liver and results in hepatosteatosis and hypobetalipoproteinemia (Zhong et al., 2010; Burnett et al., 2007; Burnett et al., 2003; Minehira et al., 2008). This highlights the ER’s essential role in lipid transport.

The stimulus-dependent secretion of proteins and lipids into circulation depends on the cell’s secretory capacity to ensure quick and sufficient signaling. Cells are equipped to respond to a stimulus with an increased secretory capacity by dynamically regulating the expression of secretory machinery genes (Feizi et al., 2017). These set of proteins include functional modules predominantly located at the ER and Golgi. These proteins recognize the mRNAs encoding proteins to be targeted to the ER, regulate their translation, translocate proteins into the ER, process the ER signal-peptide, facilitate protein folding and glycosylate proteins within the ER and traffic vesicles between the ER and Golgi (Aviram and Schuldiner, 2017; McCaughey and Stephens, 2019). Hormones, receptors, coagulation factors, extracellular matrix proteins and apolipoprotein particles are all critical clients of this machinery. Therefore, secretion machinery components serve an essential step for proteins to be processed and trafficked to the extracellular space.

But how do ER function and gene expression intersect? One evident mechanism is represented by the regulation of a handful of transcription factors that are ER-anchored and monitor the state of the ER. The most tightly linked to lipid metabolism are the Sterol regulatory element-binding proteins (SREBPs) (Shimano and Sato, 2017), notably SREBP2, which is regulated by ER membrane cholesterol content to maintain cholesterol homeostasis (Moslehi and Hamidi-Zad, 2018). Another transcription factor associated with the ER is the X-box-binding protein 1 (Xbp1) (Plumb et al., 2015), which regulates the stress response to the influx of nascent polypeptides that go beyond the ER’s protein folding capacity (Fox and Andrew, 2015). While both transcription factors regulate ER homeostasis, we know little about how cells dynamically respond to nutritional cues by coordinating transcription to impact ER homeostasis.

To address how feeding regulates nutrient and lipoprotein homeostasis *via* changes in gene expression, we took advantage of the genomic, reverse genetic and proteomic tools available in the fruit fly *Drosophila melanogaster*. We developed a fasting and refeeding paradigm to systematically profile gene activity in response to nutrients. By monitoring changes in gene expression by genome-wide profiling of gene expression and RNA polymerase II (Pol2) function, we identified an evolutionary conserved transcription factor, CrebA, which regulates cellular secretory capacity in response to feeding. We extended our analysis to show its mammalian orthologs of the Creb3L transcription factor family (Khan and Margulies, 2019) are acutely upregulated upon refeeding to coordinate the expression of genes with roles in sorting proteins into the ER and between the ER and the Golgi. We also tested whether CrebA levels alter fly feeding behavior and the levels of circulating ApoB protein. Our data reveal that this conserved family of transcription factors plays an integral role in responses to nutrition, metabolism and feeding behavior by regulating the complex biology and functions of the ER.

## RESULTS

### Fasting periods coordinate the temporal dynamics of animal feeding

To understand how long and how much flies eat during the transition from a state of hunger to satiation, we explored the dynamics of food consumption using the CAFÉ assay (Ja et al., 2007) (Figures 1A and S1A). Flies naturally exhibit circadian feeding patterns (Ro et al., 2014). They eat “meals” twice a day, once in the morning and once in the evening, but also tend to snack in between. For this reason, we limited feeding to a time-restricted schedule, and compared the amount eaten and the timing of feeding following 3, 6 or 24 hours of fasting with flies that had access to food *ad libitum*. CAFÉ revealed that the length of fasting alters how much flies eat within the first hour upon being reintroduced to food (Figure 1B). During the first hour of refeeding, flies fasted for 24 hours ate more and faster than those fasted for 3 and 6 hours. Strikingly, once fasted flies were done feeding within the first hour, they ate similar amounts as continuously-fed flies during the next 5 hours (Figure 1C). This indicates that flies “satiate” within the first hour of regaining access to food. Having established this fasting/refeeding strategy to synchronize fly feeding, we investigated the regulatory gene expression pathways that respond to feeding.

**Figure 1:**
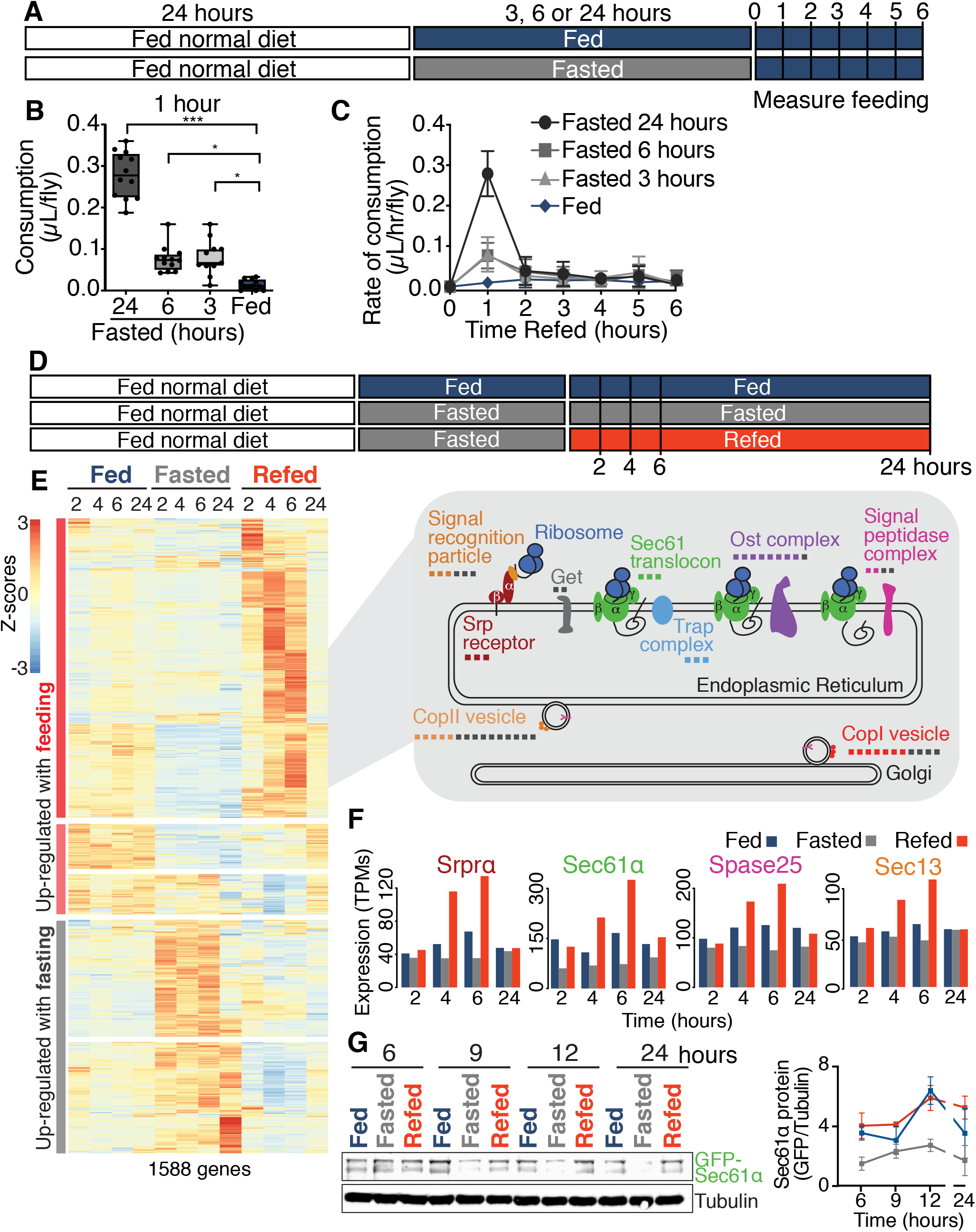
Feeding coordinates gene expression of the ER protein sorting machinery. **(A)** Scheme illustrating the experimental regiment for measuring food consumption of flies exposed to fasting or fed conditions. **(B)** Food consumed for 1 hour by female flies which are either normally fed or fasted for different amounts of time are compared. Data (n=12, except the 3 hr fasting condition n=11) are presented as box-and-whiskers plot (min to max), analyzed with Kruskal-Wallis and post-hoc Dunn’s multiple comparisons test (***p<0.0001; *p<0.05). **(C)** Rate of food consumption between flies maintained for different amounts of fasting time are compared. Data are graph as the mean ±SD (n=12, except the 3 hr fasting condition n=11). **(D)** Experimental scheme outlining the feeding regiment and sample collections for transcriptomic analysis. **(E)** Clustered heatmap of 1588 transcripts changed between pairwise comparisons of all conditions using maSigPro, clustered and visualized using pheatmap. 6 hours after refeeding, 58 out of 120 genes belonging to ER protein sorting machinery genes are enriched. In gray, a diagram shows some of the core components of the ER protein sorting machinery genes enriched in this cluster. The small squares under the name of the complexes represent the genes encoding ER protein complexes. The colored squares indicate the number of genes in each complex which have transcripts that change their levels in response to nutrition. **(F)** Barplots of the RNAseq data showing expression levels for several core components of ER sorting machinery, Signal particle receptor (Srprα), Sec61 translocon (Sec61α), Signal peptidase (Spase25) and CopI vesicle (Sec13) (n=2). **(G)** Western blot analysis with quantitation (n=3) showing the dynamics of the Sec61α during the refeeding time course, using Tubulin as a loading control.

### Feeding coordinates gene expression of the ER protein sorting machinery

To identify novel mechanisms regulating appetite, satiety and metabolism, we used expression profiling during a refeeding timecourse following an overnight fast. We focused on early timepoints after refeeding, since we were interested in dissecting the dynamics of gene regulation upon feeding to identify novel pathways regulating metabolism. To control for gene expression changes due to circadian rhythm, we compared wild-type female flies (WT) fed standard fly food, after which they were either continuously fed, fasted or fasted overnight for 16 hours followed by refeeding. mRNA was collected from heads at early time-points (2, 4 or 6 hours) and a later time-point (24 hours) and sequenced (Figure 1D). Principal component analysis of the mRNA expression data revealed a clear separation between fed and fasted, with little change within the groups throughout the timecourse (Figure S1B). In contrast, the refed sample changed during the timecourse. Indeed, the refed expression profile clearly differed from the fasted and continuously fed samples at the 2-hour time point, indicating that rapid changes in gene expression occurred within 2 hours of refeeding. Over time (24 hours) ‘re-fed’ gene expression transitioned to the ‘continuously-fed’ samples, consistent with a return of gene activity to the continuously-fed state.

We performed pairwise comparisons between fed, fasted and refed groups at each timepoint, identifying 1588 nutritionally-responsive genes (Table S1). To identify genes that might be coregulated, we used clustering analysis on the normalized levels of these transcripts during the timecourse. This revealed 621 genes with transcripts that are upregulated upon fasting and downregulated with refeeding. The upregulated “fasting genes” are enriched for proteins involved in carbohydrate and lipid metabolism (Figures 1E, S1C and S1D, Cluster 3). More than half of the nutritionally-responsive genes were upregulated with refeeding. Within the upregulated cluster, a more refined analysis of the clusters identified transcripts that were rapidly and transiently upregulated within 2-6 hours of refeeding (Figure S1C, Cluster 1). Other transcripts were upregulated more slowly, over a 24-hour time period (Figure S1C, Cluster 2), going from a low expression level in the fasted state and returning to levels seen in the continuously-fed state.

Interestingly, GO term analysis of the transiently upregulated transcripts at the 6-hour refeeding timepoint identified a cluster highly enriched for genes encoding proteins responsible for protein targeting to the ER and transporting cargo back and forth between ER and Golgi, which we will refer to as ‘ER protein sorting machinery genes’ (Figure E and S1C, Cluster 1). These genes include proteins incorporated into complexes of the Signal recognition particle (Srp), Srp receptor, Sec61 translocon, including Sec62 and Sec63, Trap complex, Signal peptidase complex, Ost complex, CopII vesicle proteins, such as Sec23 and Sec13, and CopI coat proteins (including α, β’, γ, ε and δ Cop), in addition to several p24 proteins (Figures 1E and 1F). Our data thus revealed a coordinated gene expression response to feeding of ER protein sorting machinery genes.

### The Sec61 translocon is regulated by fasting and feeding

Since the Sec61 translocon complex is the gateway into the ER for many proteins (Rapoport et al., 2017), we tested whether fasting and refeeding regulated its protein levels. We used a N-terminal GFP tagged Sec61α expressed in flies from a fosmid containing 29 kbp fragment of chromosome 2, including the Sec61α promoter and its coding region (Sarov et al., 2016). Western blotting with an anti-GFP antibody indicated that the levels of Sec61α protein follows the nutritional regulation of its mRNA transcript (Figures 1F, 1G and S1E). While fasting lowered both Sec61α mRNA and protein levels, refeeding boosted transcript levels beyond even the continuously-fed state. Further, Sec61α protein levels recovered to levels seen in continuously-fed flies. Not surprisingly, changes in Sec61α protein levels were delayed compared to transcript level changes. Protein level changes were obvious following 9 hours of refeeding, compared to 4 hours at the mRNA level. These data clearly indicate that translocon levels are impacted by nutritional state.

### Refeeding increases expression of factors required for lipoprotein particles

In addition to the core ER sorting machinery transcripts, gene transcripts required to build vesicular apolipoprotein particles were also identified in the upregulated cluster upon refeeding, together with the general ER sorting machinery. These transcripts include Tango1 (Transport and Golgi organization 1), a member of the MIA/cTAG family and a component of specialized CopII vesicles for transporting large molecules from the ER to the Golgi (Saito et al., 2009; Dreyer et al., 2019). It specializes in cargos such as collagens, preVLDL and prechylomicron lipoprotein particles (Ríos-Barrera et al., 2017). Three other transcripts also upregulated in this cluster required for the formation of VLDL and the transport of ApoB in cell culture included the Microsomal triglyceride transfer protein (Mtp), its interaction partner, the Protein disulphide isomerase (Pdi) (Wetterau et al., 1990) and Surfeit 4 (Surf4) (Wang et al., 2020) (Figure S1F).

Interestingly, the levels of transcripts encoding components that regulate the incorporation of proteins into the ER membrane, such as the guided entry of the tail-anchored proteins (GET pathway) (Schuldiner et al., 2008) and the mammalian Tmco1-dependent pathway responsible for the insertion of multipass transmembrane proteins into the ER membrane (Shurtleff et al., 2018) were not nutritionally regulated. In addition, other ER “stress” pathways did not change upon flies refeeding. For example, neither Ire1 of the Unfolded Protein Response (UPR) pathway (Shen et al., 2001; Yoshida et al., 2001), nor components of the ER-associated protein degradation (ERAD) pathway (Lopata et al., 2020). To summarize, fasting and refeeding specifically regulated transcripts involved in protein sorting into the ER, including the Sec61 translocon, but not all components associated with ER function (Figure S1F and Table S1).

### The *Drosophila* transcription factor CrebA is regulated by feeding

To investigate the mechanisms that drive changes in the expression of ER protein sorting machinery genes upon feeding, we searched for more immediate changes in our RNAseq data. The coordinated feeding- and time-dependent regulation of multiple ER protein sorting machinery genes hinted toward a single regulator, whose own expression levels might have changed at earlier timepoints in our timecourse. Interestingly, we found that the transcript encoding the basic leucine zipper transcription factor, CrebA, responded quickly to fasting and feeding (Figure 2A). The fact that CrebA has been shown to regulate ER sorting machinery genes during early fly development (Fox et al., 2010) made it a candidate for regulating these genes in response to feeding. RTqPCR assays validated CrebA transcript changes observed in our RNAseq data, indicating that CrebA mRNA was acutely upregulated within 2 hours of refeeding (Figure 2B) and over time decreased back to continuously-fed levels. CrebA protein levels followed similarly stark dynamics of CrebA mRNA regulation upon fasting and feeding (Figure 2C). Fasting abrogated CrebA mRNA and protein levels, while refeeding rapidly induced both CrebA mRNA and protein. Our data point to CrebA as a potential regulator of coordinated gene expression changes of the ER protein sorting machinery in response to nutrients.

**Figure 2:**
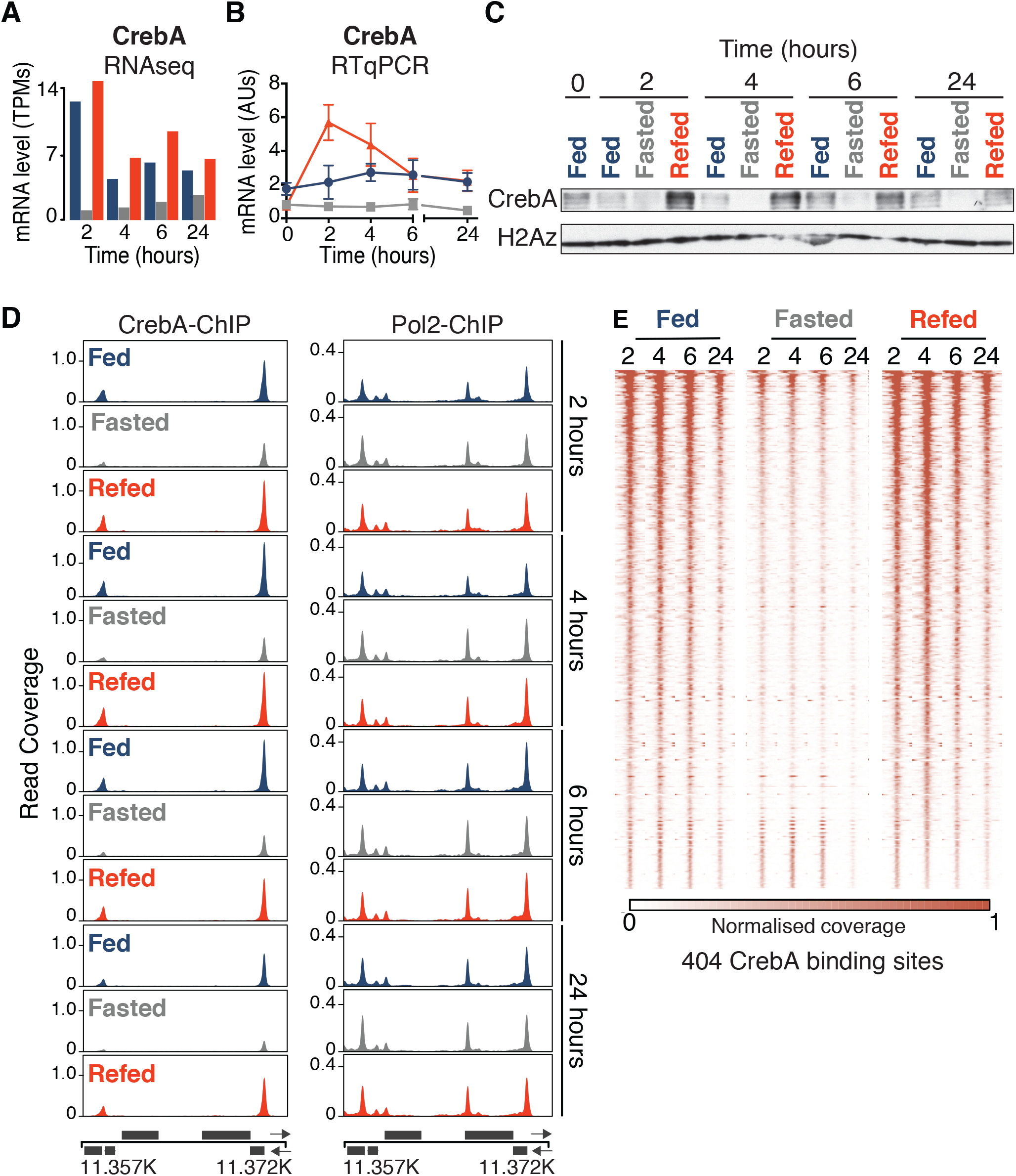
The transcription factor CrebA is regulated by feeding. **(A)** CrebA expression levels during the refeeding timecourse graphed using the RNAseq data (n=2) and **(B)** assessed by RTqPCR (normalized to H2A.Z). RTqPCR data are means ±SD for n=3-5. **(C)** Representative western blots showing CrebA and H2A.Z (loading control) protein levels. **(D)** Representative ChIPseq profiles of CrebA (left) and Pol2 (right) along the refeeding time-course. Region plotted is 2R:11.357.693-11.372.933. **(E)** A heatmap showing bound CrebA levels +/−750 bp centered over 404 strong CrebA binding sites as identified by ChIPseq analysis. Regions are presented in decreasing order based on the signal at refed 4-hours.

To probe this hypothesis, we investigated whether CrebA activity is regulated by feeding using genome-wide chromatin-immunoprecipitation assays (ChIPseq) of CrebA. First, we validated a polyclonal CrebA antibody (Figures S2A, S2B and S2C). Next, we assayed CrebA-binding genome-wide and found that CrebA bound its target genes in a manner that depended highly on the feeding status and time (Figures 2D, 2E and S2D, Table S2). Most CrebA sites mapped to annotated promoters, transcription start sites (TSS) or within the 1^st^ exon of the gene (Figure S2E). Overall, CrebA binding to 404 sites across the whole genome was increased with refeeding, with the highest occupancy occurring at the 4-hour timepoint. These CrebA ChIPseq changes were not due to differences in chromatin preparations, as indicated from a control ChIPseq from the same chromatin for RNA Polymerase II (Pol2). From this control, we observed that global paused Pol2 peaks did not change on the promoters of genes between different nutritional conditions (Figure 2D, S2D and S2G, Table S2). This indicates that CrebA activity is regulated by feeding. Additionally, an unbiased *de novo* search for sequence motifs in the CrebA-bound regions identified a robust enrichment for the known CrebA consensus sequence, further validating our ChIPseq results (Figure S2F) (Abrams and Andrew, 2005; Nitta et al., 2015; Johnson et al., 2020). Overall, our results highlight a rapid response of CrebA binding to its target genes upon refeeding.

### CrebA regulates ER protein sorting machinery gene transcription upon feeding

Since both CrebA expression levels and its chromatin binding are clearly regulated by nutritional status, we asked whether CrebA could act as the upstream transcriptional regulator of the ER protein sorting machinery genes in response to feeding. We found that one of the main components of the secretory pathway, the Sec61 translocon, was strongly bound by CrebA in a nutrition-dependent manner (Figure 3A and D). Upon annotating all CrebA-bound regions to the nearest genes, we found 518 genes with a CrebA-binding site within 1500 bp of the core promoter region of these genes. Remarkably, GO term analysis of this gene set revealed a strong enrichment for the ER protein sorting machinery genes that we found to be transiently upregulated after refeeding (Figure 3B). ChIPseq analysis showed that many ER protein sorting genes were CrebA targets. Interestingly, comparison of CrebA targets with the genes identified as nutritionally-responsive showed that not all CrebA-targeted genes encode for transcripts that changed with nutritional cues (Figure 3C and S3C). However, of the 149 CrebA target genes whose transcripts did change with nutrition, 48 of these belonged to nutritionally-regulated ER protein sorting genes, including Tango1, Surf4 and Pdi, the binding partner of MTP, both of which showed CrebA binding at their promoters. In addition, all of the 149 nutritionally-regulated CrebA target genes were shown to have upregulated transcripts upon refeeding. This reveals CrebA as the master regulator of cellular secretory capacity in response to nutritional cues.

**Figure 3:**
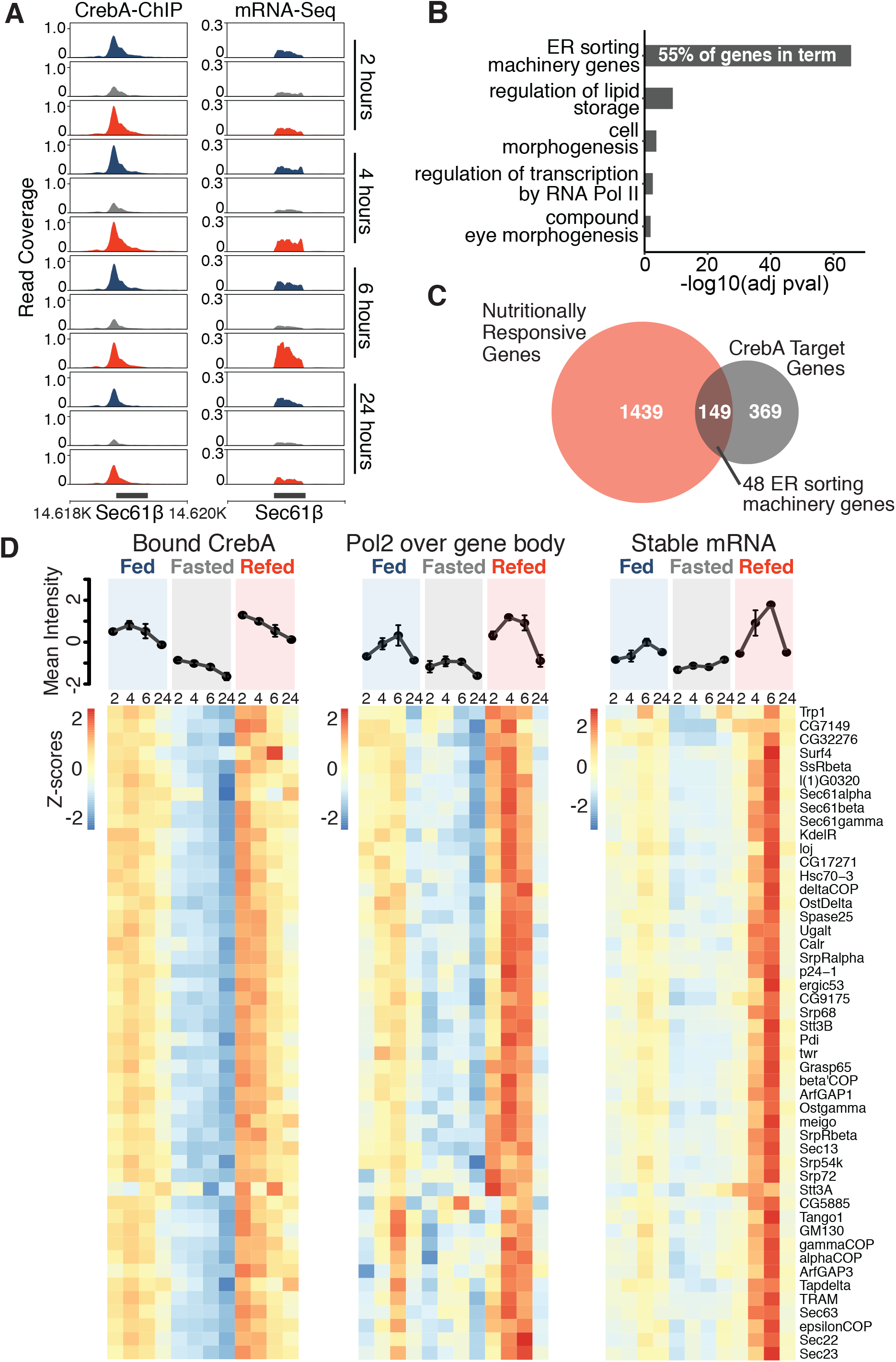
CrebA regulates the transcription of ER protein sorting machinery genes in response to feeding. **(A)** Representative CrebA ChIPseq (left) and RNAseq (right) profiles from fly heads over Sec61β - a core component of the ER sorting machinery. Region plotted is 2R:14.617.400-14.620.853. **(B)** Functional enrichment analysis of 518 identified CrebA targets using Fischer’s-Exact test. The most significant GO-terms are shown with FDR <0.05. **(C)** A Venn diagram indicating the overlap between 1588 nutritionally-responsive genes identified using RNAseq and 518 CrebA target genes. **(D)** A heatmap showing individual z-scores plotted for the 48 ER protein sorting machinery genes for CrebA binding, Pol2 signal over the gene body and mRNA levels. All heatmaps are row clustered based on hierarchical clustering performed on Pol2 signals. Mean z-scores for all genes are graphed on top.

### CrebA acts as the metabolic switch that promotes RNA polymerase II elongation

To compare how CrebA activation relates to transcription, we plotted signals for bound CrebA, Pol2 and mRNA levels over the 48 ER protein sorting machinery genes (Figure 3D). Notably, we observed a clear increase of CrebA binding at the promotor of these genes within 2 hours after eating. This was followed by an increase in Pol2 elongating over these genes, peaking at 4 hours (Figure 3D), after which mRNA levels increased collectively, peaking at 6 hours. Next, we used Pol2 ChIPseq to determine if these CrebA targets were regulated by Pol2 promoter recruitment or Pol2 elongation. Pol2 levels over the gene body (minus the TSS) over the timecourse (Figure S3A) showed more dramatic changes than changes in Pol2 over the TSS (Figure S3B). The pattern of Pol2 increase over the body of these genes in the refed state suggests that Pol2 activity at these genes is regulated by elongation, rather than recruitment of Pol2 to the promoter. This suggests that CrebA likely activates transcription by facilitating Pol2 promoter escape. The extensive and robust feeding-and time-dependent transcriptional regulation of ER protein sorting genes by CrebA supports our hypothesis that CrebA is a key metabolic switch that upregulates a core set of ER protein sorting genes.

### Nutritional state regulates changes in the hemolymph proteome

After establishing the regulatory mechanism coordinating the expression of ER protein sorting genes, we asked how changing the levels of the Sec61 translocon and other components of the ER sorting machinery would impact ER function. We hypothesized that changes to the levels the ER sorting machinery upon feeding could alter the ability of cells to secrete proteins. To orthogonally complement our transcriptomic refeeding timecourses, we used mass spectrometry (MS) to quantify changes in protein secretion depending on nutritional state. We thus harvested the fly hemolymph from fed, fasted and refed flies, and identified the proteins using liquid chromatography-based MS (LC-MS) (Figure S4A). Principal component analysis showed clear separation of fed and fasted groups in the first component PC1 (Figure 4A). Over time, refeeding led the proteome to change from the fasted condition to resemble the continuously-fed proteome. From the total of 1878 hemolymph proteins identified by LC-MS, 405 proteins levels changed with nutritional status (Table S3). Because the hemolymph also contains cells, referred to as hemocytes, which have a role in innate immunity and wound healing (Lemaitre and Hoffmann, 2007; Honti et al., 2014), we checked whether our dataset included cellular proteins. The hemolymph MS analyses revealed an enrichment of ER-targeted proteins, with 66% of the proteins detected in the hemolymph carrying a signal peptide (Figure S4B). The proteins with a signal peptide were enriched for extracellularly-localized proteins (Figure S2C). Clustering of these proteins with a signal peptide based on their levels during our fasting/refeeding timecourses identified groups of proteins that were upregulated with refeeding (Figure 4B) and group of proteins upregulated during fasting. In addition, there is a cluster of proteins that showed only transient changes after refeeding. Included in the upregulated with refeeding cluster were secreted proteins including Closca (Clos), Fs(1)N and Fs(1)M3, which are involved in Torso signaling (Ventura et al., 2010) (Figure S4D). The transient, but coordinated changes in the levels of these hemolymph proteins may identify a role for Torso signaling in response to nutrition.

**Figure 4:**
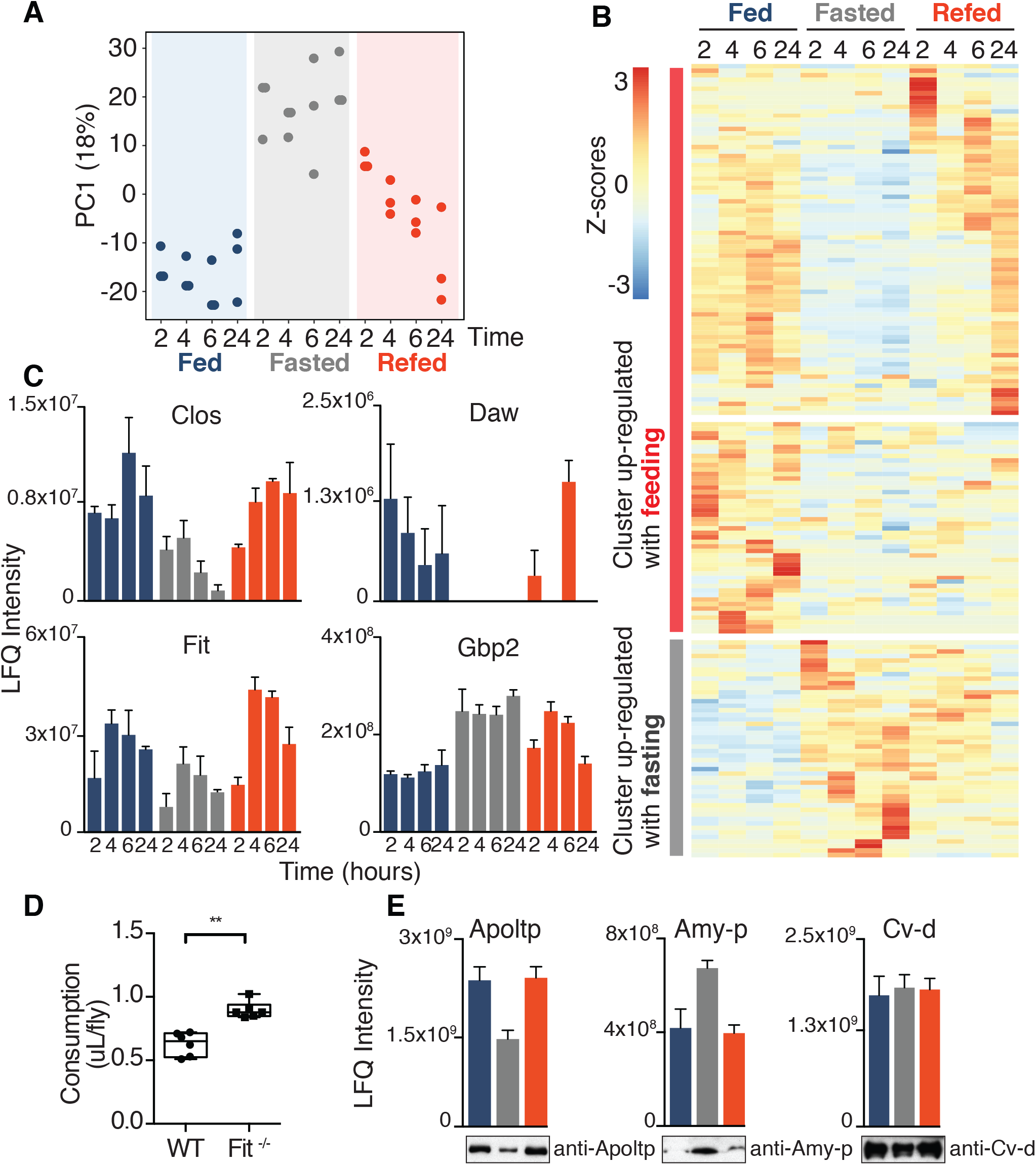
Hemolymph proteome changes with nutritional state. **(A)** Principal component analysis showing the first component plotted against condition and time of 1878 detected proteins from the hemolymph MS. **(B)** A heatmap showing 405 differentially present proteins containing a signal peptide in the hemolymph. Z-scores plotted for n=3. **(C)** Barplots of MS data showing protein expression of Fit, Daw, Clos and Gbp2 proteins along the refeeding timecourse. LFQ normalized intensities plotted for n=3 ±SEM. **(D)** Fit mutant (Sun et al., 2017) had higher food intake compared with wild-type flies in CAFÉ. Data (n=6) are presented as box-and-whiskers plot (min to max) and analyzed with Mann-Whitney test (p=0.002). **(E)** Barplots showing levels of Apoltp, Amy-p and Cv-d from the MS hemolymph data and complementary western blots of hemolymph harvested 24 hours after refeeding. LFQ normalized intensities plotted for n=3 ±SEM.

In addition, our hemolymph data identified feeding-dependent hormones that are transiently increased upon refeeding. These included Dawdle (Daw), the sugar signaling TGF-β ligand (Chng et al., 2014; Chng et al., 2017) and Female-specific independent of transformer (Fit) (Figure 4C), a satiety hormone (Sun et al., 2017). Consistent with its role regulating satiety, *Fit* mutants feed more than wild-type flies (Figure 4D). Growth-blocking peptide 2 (Gbp2), a fat body peptide that signals reduced growth rates (Koyama et al., 2020), was upregulated in the fasted state. Other proteins with altered expression in the hemolymph included proteins involved in lipid metabolism. Several lipases, including YP1, YP2, YP3 and CG5162, were all upregulated in the hemolymph upon fasting (Figure S4D). In later refeeding timepoints, other proteins with roles in lipid metabolism were upregulated, such as Saposin-related (Sap-r), involved in sphingolipid metabolism (Sellin et al., 2017), Glutactin, a member of the cholinesterase family (Olson et al., 1990), Nieman-PickC2 (Npc2), involved in cholesterol metabolism (Huang et al., 2007), and the *Drosophila* ApoB ortholog, Apoltp, which is implicated in building hemolymph lipoprotein particles with lipids from the gut (Palm et al., 2012).

Next, we validated changes in hemolymph protein levels identified by LC-MS by western blots. This showed that Apoltp levels decreased in the hemolymph upon fasting and recovered to the continuously-fed state upon refeeding (Figure 4E). In contrast, Amylase proximal (Amy-p) was upregulated in the fasted state in the same hemolymph preparations, while Crossveinless d (Cv-d) levels remained (Figure 4E), validating our LC-MS data. Altogether, MS revealed nutrition-dependent changes in the hemolymph levels of hormones, signaling proteins and other proteins involved in metabolism. Likely, many of these changes are the direct result of fasting and refeeding changing the activity of the ER protein sorting machinery, thus regulating the entry of these proteins into the ER and impacting their secretion. We suggest that CrebA indirectly regulates the changes in the levels of these hemolymph proteins by throttling the expression of the ER sorting machinery in response to feeding.

### Transient CrebA overexpression suppresses animal feeding

Since satiated flies showed higher CrebA expression levels, we hypothesized that overexpressing CrebA could mimic the fed state and flies might thus eat less. The overexpression experiment was performed by transiently expressing CrebA using the bipartite Hsp70-Gal4 system (Brand and Perrimon, 1993). Flies were fed on standard food for 24 hours before heat shock. After heat shock, they recovered overnight for 16 hours (Figure 5A). Flies were then fasted for 6 hours to induce hunger and feeding was measured using CAFÉ (Figure S1A). Western blots probing for CrebA showed that CrebA was overexpressed during the entire experiment (Figure S5A). Interestingly, while wild-type, genetic control and heat shock control flies showed no changes in food consumption, flies with overexpressed CrebA decreased feeding (Figures 5B). Since flies must climb to eat in the CAFÉ assay, we checked whether overexpressing CrebA hindered fly mobility. Using similar controls as in the CAFÉ assay, CrebA overexpression did not alter fly mobility (Figures 5C and S5B). Our results reveal that increased CrebA levels suppress food consumption.

**Figure 5:**
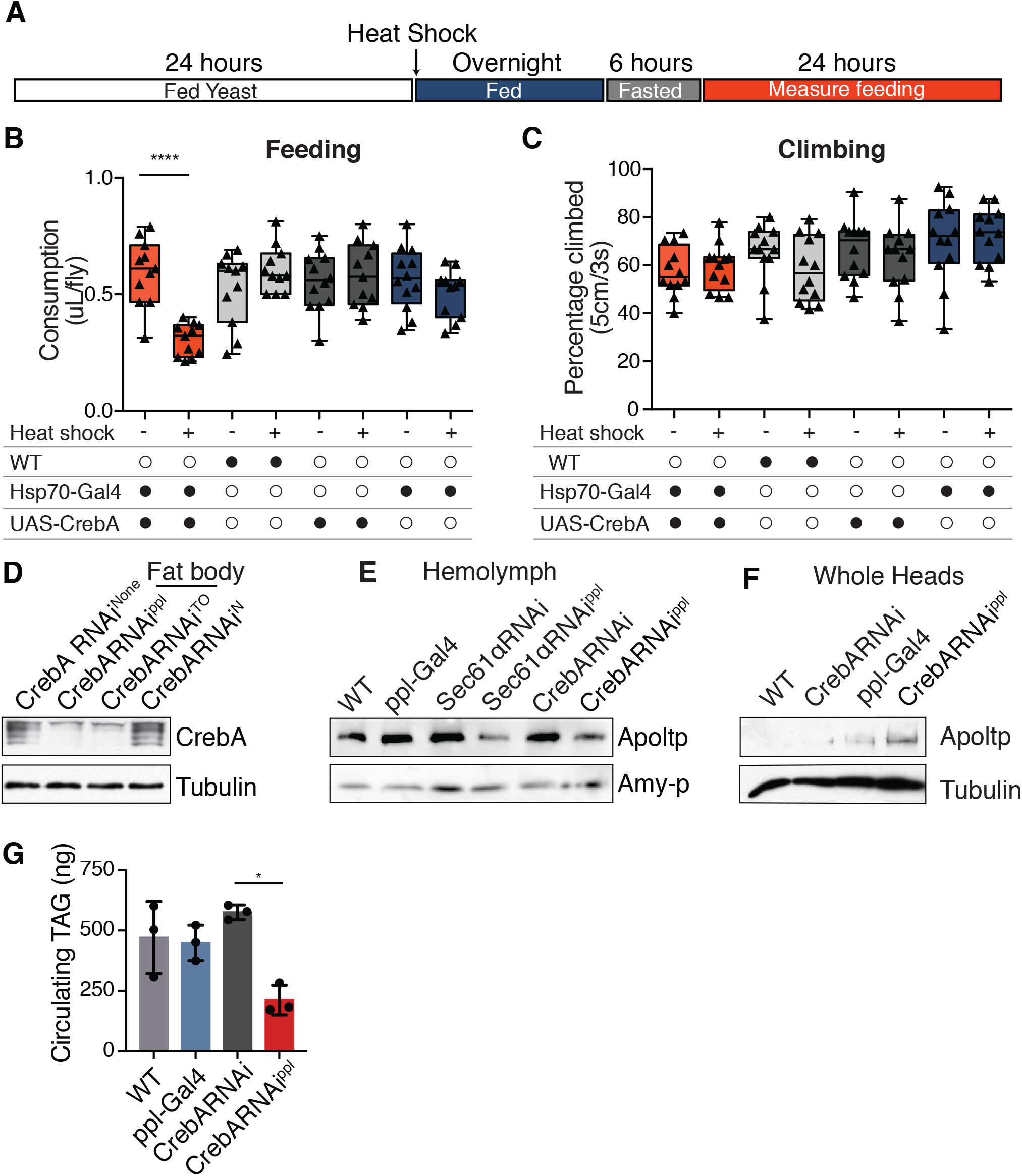
CrebA levels alter feeding and secreted lipoprotein in the hemolymph. **(A)** Schematic outlining the treatment of flies until measuring their food consumption. Flies were heat-shocked for 20 minutes at 36°C. **(B)** Food consumption of flies over 24 hours using the CAFE assay upon overexpression of CrebA. Data (n=12) are presented as box-and-whiskers plot (min to max), analyzed with Kruskal-Wallis and post-hoc Dunn’s multiple comparisons test (*p<0.03,** p<0.002; ***p<0.0002). **(C)** Fly climbing performance measured using the RING assay (Gargano et al., 2005). Data (n=12) are presented as box-and-whiskers plot (min to max). **(D)** Representative western blot showing CrebA and Tubulin protein (control) levels upon the knockdown of CrebA in different cell-types. Fat body drivers are ppl and TO. The neuronal driver is nSyb (N). **(E)** Representative western blot showing levels of Apoltp and Apolpp protein levels in hemolymph from 20 flies upon knockdown of Sec61α and CrebA in fat body cells. **(F)** Representative western blot showing levels of Apoltp and Tubulin levels in heads the knockdown of CrebA. **(G)** The levels of circulating TAG from 25 flies upon CrebA knock down in fat body cells (n=3).

### The fat body is the prevalent cell type for feeding-regulated CrebA

Identifying the cell type(-s) where CrebA is nutritionally regulated, provides insights into the physiological role of changing cellular secretory capacity in response to feeding. Thus far, our transcriptomics, qPCR and western blot were all performed on fly heads. To dissect the cell-type-specific roles of CrebA in mediating an organism’s systemic response to nutritional changes, we focused on the cell types found in the *Drosophila* head, such as neurons, glia, muscle cells and the fat body, which has functions analogous to those of vertebrate hepatocytes and adipocytes. We first knocked down CrebA expression in neurons, because they constitute the most abundant cell type in the head, and in the fat body, because of their important metabolic function. Using a UAS CrebA shRNAi fly line (Dietzl et al., 2007), we knocked down CrebA in fat bodies using either takeout-Gal4 (TO) (Dauwalder et al., 2002) or ppl-Gal4 (ppl) (Zinke et al., 1999) drivers. Under these conditions, no CrebA was detected in the fly head 2 hours after refeeding initiated (Figure 5D). In contrast, CrebA was detected in fed flies using the CrebA shRNAi alone or when the driver nSyb-Gal4 (N) was used to knock down CrebA in neurons. These knockdown experiments revealed that the majority of CrebA upregulated upon refeeding was expressed in fat body cells.

### CrebA regulates lipoprotein secretion into the hemolymph

We hypothesized that CrebA plays an important role in regulating the secretory capacity of the fat body and could therefore impact metabolite distribution and interorgan communication of the metabolic state. Indeed, insect fat body cells secret proteins involved in diverse processes, including proteins that regulate lipid transport throughout the body (Palm et al., 2012). *Drosophila*‘s two ApoB-family lipoproteins Apolpp and Apoltp are both produced in the fat body. Apoltp loads lipids from the gut into Apolpp particles (Palm et al., 2012), but may also signal to the central nervous system (CNS) (Brankatschk et al., 2014). Our hemolymph proteome did not identify nutrient-regulated changes in Apolpp protein. However, Apoltp protein decreased upon fasting and returned to fed levels upon refeeding (Figures 4E). We therefore further dissected its regulation by CrebA and transport to the ER. Apoltp has an ER N-terminal signaling peptide, so Apoltp likely requires the Sec61 translocon to be secreted into the hemolymph. To test this formally, we knocked down Sec61α in the fat body. Hemolymph collected from fat body-Sec61α-knockdown flies exhibited lower levels of Apoltp, compared with wild-type and other genetic controls (Figure 5E). We next addressed whether CrebA regulates Apoltp hemolymph levels. Interestingly, Apoltp levels were decreased in the hemolymph (Figure 5E), but not in total head extracts when CrebA was selectively knocked down in the fat body (Figure 5F). In fact, in total head extracts, the amount of Apoltp increased, suggesting that secretion and not expression of Apoltp was altered by CrebA activity. The fat body CrebA knockdown flies also have reduced circulating triacylglycerol (TAG) (Figure 5G), corroborating the role of CrebA in lipid metabolism. Our data support the hypothesis that CrebA regulates the secretion of proteins from the fat body by regulating the Sec61 translocon.

### Feeding regulates the expression of mammalian CrebA orthologues

Many metabolic and nutrient-regulated hormonal signaling pathways are conserved between *Drosophila* and mammals. We therefore asked whether mammalian CrebA orthologs are nutritionally regulated. Mammals have five CrebA orthologs (Khan and Margulies, 2019) (Figure 6A). Creb3L1 and Creb3L2 recognize the same DNA sequence. The DNA binding region of CrebA and Creb3L2 are identical and the other mammalian orthologs have very similar basic regions. In addition, if we compare our *de novo* search for CrebA binding sites from our ChIPseq (Figure S3E) with published ChIPseq for Creb3L1 and Creb3L2 (Consortium, 2012; Davis et al., 2018; Khetchoumian et al., 2019), we found the same DNA motif for all 3 family members (Figure S6C). Further, mammalian Creb3L family members are able to activate expression of the same genes as CrebA targets when expressed in flies (Fox et al., 2010; Barbosa et al., 2013). In general, CrebA orthologs have roles in biological processes that require secretion, such as in bone/cartilage development and liver function (Khan and Margulies, 2019). Since we found CrebA to be nutritionally regulated in the fat body, whose roles are analogous to the mammalian liver, we targeted the function of Creb3L family members in the liver.

**Figure 6:**
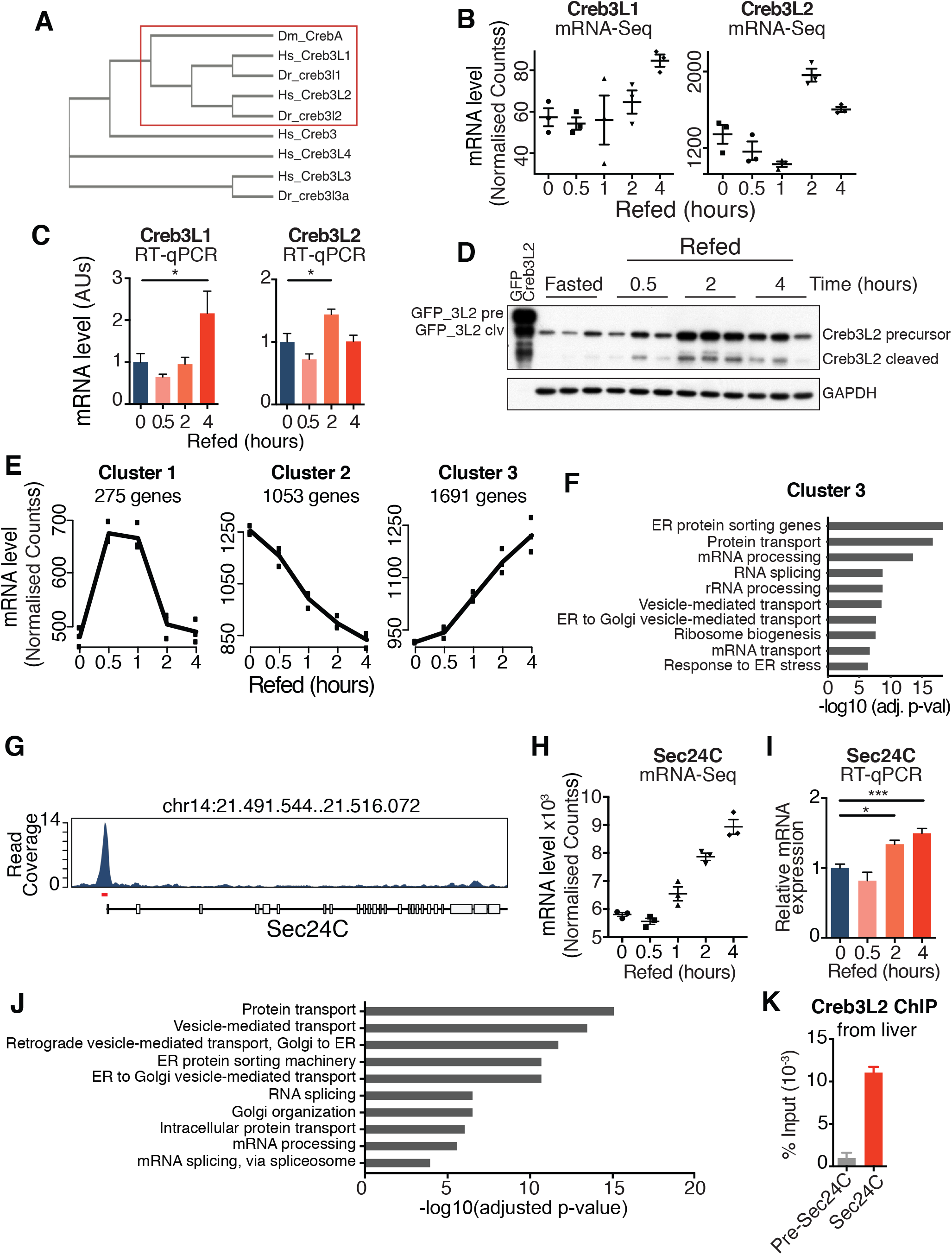
Mammalian homologs of CrebA regulate the ER protein sorting machinery genes in response to feeding. **(A)** Phylogenetic cladogram of CrebA mammalian and zebra fish homologs based on a sequence blast. Diagram was made using clustal Omega. **(B)** Creb3L1 and Creb3L2 mRNA expression levels in mRNAseq data from mice liver upon refeeding. Data are means ±SEM for n=3 biological replicates (Brandt et al., 2018). **(C)** RTqPCR measured mRNA levels of Creb3L1 and Creb3L2, normalized to a housekeeping TBP gene, in mice liver upon refeeding. Data (n=6 biological replicates) analyzed with one-way ANOVA with Tukey correction (*p<0.05). **(D)** Representative western blot showing levels of Creb3L2 in mice liver along the refeeding time-course (3 different animals for each time point). GAPDH shown as loading control. **(E)** Clustering analysis of the RNAseq dataset (Brandt et al., 2018) from refed mice liver along the timecourse performed with maSeqPro. **(F)** Functional enrichment of 1691 genes in cluster 3 using Fischer’s Exact test. **(G)** Coverage tracks showing Creb3L2 binding one of the secretory machinery genes, Sec24C (Khetchoumian et al., 2019). **(H)** mRNA levels of Sec24C upon refeeding (Brandt et al., 2018), n=3. **(I)** mRNA levels of Sec24C upon refeeding using RTqPCR, n=6. Data plotted min to max and analyzed using one-way ANOVA with Tukey’s test compared with all groups (***p<0.001 and *p<0.05). **(J)** Functional enrichment of all Creb3L2 target genes using Fischer’s Exact test using the ChIPseq data from Khetchoumian et al., 2019. **(K)** ChIP of Creb3L2 from liver chromatin using primers over the Creb3L2 peak in marked in red panel 6G. The control primers are located 3kb upstream of the Sec24 promoter.

To investigate which Creb3L family members might be regulated by feeding in the mouse liver, we mined RNAseq data from the liver of mice refed 0, 0.5, 1, 2 and 4 hours on a chow diet (Brandt et al., 2018). Interestingly, both Creb3L1 and Creb3L2 were upregulated upon refeeding (Figure 6B). To validate these genome-wide results, we collected liver samples harvested from mice refed for 0.5, 2 and 4 hours. RTqPCR on these samples recapitulated our observed *Drosophila* CrebA regulation, with a similar acute upregulation of Creb3L1/L2 expression upon refeeding (Figure 6C). Further, western blots with Creb3L2 antibodies indicated that the protein levels for this isoform changed transiently in the liver (Figure 6D), recapitulating CrebA regulation by nutrients in the fly. Our data indicate that a mammalian CrebA ortholog is also regulated by nutrients. Interestingly, unlike fly CrebA, mammalian Creb3L isoforms have a C-terminal ER anchoring domain (Omori et al., 2002). In response to nutrients, this cleavage of Creb3L2 is not rate-limiting, as the cleaved active N-terminus is also upregulated with nutrition. The ratio of cleaved to uncleaved Creb3L2, for example, remains the same at the different times of the timecourse (Figure 6D, S6A and S6B). The kinetics of the transcriptionally active, cleaved form – combined with the transcript levels – suggest that feeding regulates Creb3L2 at the transcript level, similar to fly CrebA.

### Mammalian CrebA homologs regulate ER protein sorting machinery genes

Next, we investigated whether ER protein sorting machinery genes are regulated by nutrients in mammals. While these genes are highly conserved in eukaryotes (Lang et al., 2017), it is not clear whether their regulation by nutrients is conserved. Mining published data from liver (Brandt et al., 2018), unsupervised clustering of mRNAseq results identified 1691 transcripts that were upregulated along the timecourse after refeeding (Figure 6E, Cluster 3). Notably, upon functional enrichment analysis, ER sorting machinery genes were highly enriched in this cluster, suggesting coregulation upon refeeding (Figure 6F). Importantly, as we observed in *Drosophila*, the mouse liver data indicated a conservation of Creb3L transcription factor and secretory machinery pathway function upon refeeding. Consistent with our findings in *Drosophila*, we therefore postulated that the coordinated response of the early secretory machinery genes is mediated by Creb3L family members.

To test this possibility, we used published ChIPseq data of Creb3L family members to test whether mammalian Creb3L isoforms target the secretory machinery genes at the chromatin level. Analyses of a Creb3L2 ChIPseq dataset from a mouse pituitary cell line (Consortium, 2012; Davis et al., 2018; Khetchoumian et al., 2019) identified 1818 targets of Creb3L2 that were highly enriched for early ER sorting genes (Figures 6J). Sec24C, a core of the CopII vesicle, was one of the strongly bound targets of Creb3L2 (Figure 6G), indicating that Sec24C is a target of mammalian Creb3L. Indeed, Sec24C mRNA levels were upregulated in the liver upon refeeding (Figure 6H), which we validated using RTqPCR (Figure 6I). In fact, using ChIP of Creb3L2 in liver, we observed that Creb3L2 bound specifically to the promoter of Sec24C (Figure 6K). Taken together, our results suggest that Creb3L family members share a conserved function in flies and mammals as metabolic switches that are acutely turned on upon (re-) feeding and – in turn – coordinate the secretory capacity of cells.

## DISCUSSION

Feeding and fasting induce systemic changes to an organism’s physiology, from regulating hunger to satiation, nutrient absorption, distribution and storage. Through its secretory capacity, the ER connects cellular responses with organism-wide effects, placing it at the crossroads of cell stress and metabolism (Koerner et al., 2019). Most appetitive hormones, for example, pass through the ER to be secreted into the bloodstream. The ER is also the location for enzymatic reactions in gluconeogenesis and lipid metabolism, regulating a cell’s ability to organize lipids into droplets for cellular storage or into apolipoprotein particles destined for export. Many secreted proteins, as well as ER- and Golgi-localized proteins, depend on the Sec61 translocon for their entry into the ER. Changing the levels of this translocon and the supporting ER sorting machinery thus affects the ER protein repertoire, which – in turn – changes the storage and secretion of proteins and lipids. Alterations in the repertoire of these proteins also contribute to ER stress and the lipid accumulation, as is often observed in metabolic diseases (Molenaar et al., 2020).

### CrebA homologs are key nutritional boosters of cellular secretory capacity

Here, we have identified a conserved transcriptional mechanism that links nutritional cues to CrebA/Creb3L transcription factor-mediated changes in the repertoire of ER proteins that sort proteins to the Golgi. Our analysis allows us to propose that this mechanism is a key regulatory node which increases the cellular secretory capacity in response to feeding, allowing organisms to respond to and better deal with changing nutrient levels. By transcriptionally upregulating the ER sorting machinery to target ER-dependent proteins, such as apolipoproteins, cells are likely to avoid ER stress. It is known, for example, that cells lacking Tango1, a CopII vesicle component required for transporting large cargos from the ER to the Golgi, activate the UPR pathway (Ríos-Barrera et al., 2017). Interestingly, our data revealed that Tango1 and other ER components were upregulated upon refeeding and that this was mediated by CrebA activation. Our data revealed that CrebA and its mammalian orthologs modulate ER function and thus critical in mediating the increased secretory capacity requirements following feeding.

### CrebA may regulate anorexigenic signals

We show that feeding increased CrebA target gene expression, which impacted the secretion of proteins. This suggests that changes in CrebA activity may impact behavior by altering the repertoire of secreted proteins, including hormones. Interestingly, we observed that increasing the expression of CrebA was sufficient to suppress feeding (Fig. 5B) and that this occurred without impacting animal movement. Although we have not addressed how CrebA suppresses food intake, we suspect that this repression of feeding could be due to alterations in the levels of anorexic signals, such as Fit (Sun et al., 2017). Indeed, our hemolymph proteomics data revealed Fit as one of several, transiently-increased hormones. We also show that Fit mutants consume more food (Figure S4E) consistent with previous findings (Sun et al., 2017). Other peptides known to regulate feeding behaviors, such as sNPF (Lee et al., 2004), Hugin (Melcher and Pankratz, 2005), CCHa2 (Ren et al., 2015), dilps and adipokinetic hormone (Kim and Rulifson, 2004; Lee and Park, 2004; Gáliková et al., 2015) were not detected in our hemolymph proteome and thus changes in concentration could not be quantitated. Targeted MS approaches may be necessary to probe for these proteins.

Beyond hormones, our study revealed changes in Apoltp hemolymph levels upon feeding. Importantly, the Apoltp hemolymph levels, but not expression levels, were regulated by CrebA. We propose that CrebA indirectly impacts hemolymph Apoltp levels by regulating the ER protein sorting machinery. In turn, Apoltp may mediate CrebA-regulated feeding. Supporting this hypothesis, LpR1, a receptor binding to apolipoproteins (Rodríguez-Vázquez et al., 2015), regulates nutrient-driven food-seeking behavior (Huang et al., 2020). Apoltp may also directly signal to the brain. Intriguingly, published data show that Apoltp secreted from the fat body localizes to the CNS in a nutrient-dependent manner (Brankatschk et al., 2014). Moreover, apolipoprotein particles may have anorexigenic roles in regulating feeding in mammals, too. Indeed, apolipoprotein receptor LRP1 forebrain knockout mice gain weight and have increased food intake (Liu et al., 2011). Further, intravenous infusion of ApoAIV reduces food intake in rats (Fujimoto et al., 1992; Wang et al., 2013). Clearly, CrebA’s ability to regulate circulating Apoltp levels and mediate food intake *via* lipoprotein complexes reveals an interesting physiological and health-relevant function of this transcription factor. Future work will elucidate the precise signaling mechanism(-s) that links nutrients to feeding behavior *via* the molecular regulation of the metabolic switch CrebA and of its orthologs.

### Nutrient regulation of mammalian Creb3L in the liver

In mammals, the transcriptional pathways discussed here are more sophisticated, since there are 5 CrebA orthologs. In this study, we found that Creb3L1 and Creb3L2 levels increased upon feeding in the mouse liver. This increase in expression with feeding is in contrast to the expression levels of Creb3L3 (CrebH), whose levels decreased with feeding (Khan and Margulies, 2019). This opens up the possibility for crosstalk between the different Creb3L isoforms. Since they may form heterodimers, which isoform is present could affect transcriptional activity. However, it also possible that these changes in individual Creb3L transcription factors occur in different cell types within the liver or that they have different roles. For Creb3L3, for example, most research has focused on its direct transcriptional function on the secreted ApoAIV (Xu et al., 2014) and other genes involved in lipid metabolism (Zhang et al., 2012; Xu et al., 2015), but to date not on the ER protein sorting machinery. However, here we show that Creb3L2 binds to the promoter of a CopII component in the liver, complementing genome-wide data and pointing to the ER protein sorting machinery as the major target of Creb3L2 function (Fox et al., 2010). Our data constitute the first report showing that Creb3L1 and Creb3L2 are regulated by nutritional cues in the liver. Our results suggest that Creb3L1 and Creb3L2 will have similar roles to the fly transcription factor CrebA in regulating ER function upon nutritional cues and affecting lipid metabolism.

### The transcriptional regulation of Creb3L transcription factors

Creb3L-family transcription factors regulate the ER protein sorting machinery genes under distinct biological circumstances. Our data reveal that they regulate the ER sorting machinery in response to nutritional state. We showed that the transcripts encoding these conserved transcription factors are upregulated with feeding. While we detected changes in Creb3L2 protein levels, likely the main level of regulation occurs at the transcriptional or translational level. In the future, it will be important to establish how CrebA and other Creb3L family members are regulated by nutritional cues.

What role could this nutrient-regulated transcriptional control of ER protein sorting machinery genes play? We suspect that the feeding-mediated upregulation of the ER sorting machinery genes may avoid ER stress. There are hints of the interplay between Creb3L family members and ER stress *via* the transcription factor Xbp1, which regulates the response to ER stress *via* the UPR pathway (Liu et al., 2019). Xbp1 appears to regulate Creb3L family members by binding to *Creb3L1* and *Creb3L2* promoters in response to nutritional state in the liver (Liu et al., 2019). In addition, Xbp1 appears to transcriptionally regulate some of the same ER protein sorting machinery genes as Creb3L family members (Liu et al., 2019). We suggest that nutritional input increases the translation of ER-targeted proteins, which transiently activate Xbp1 to promote expression of Creb3L2/CrebA. In turn, this increases the expression of the ER protein sorting machinery. These data suggest that Xbp1 may be key to regulating CrebA activity in response to feeding.

Accumulating evidence also points to feedback pathways between CrebA family members and the transcription factor Xbp1. There is evidence that Creb3L-family members transcriptionally regulate the *Xbp1* gene (Johnson et al., 2020). In flies, ChIPseq data and transcriptional reporters in embryos indicate that *Xbp1* transcription is directly activated by CrebA (Johnson et al., 2020). Further, we see CrebA bound to the *Xbp1* promoter in adult heads (Figure S3C). This indicates that CrebA has a direct role in activating *Xbp1* transcription. However, CrebA also seems to act as a repressor of *Xbp1* expression in response to bacterial infection (Troha et al., 2018), suggesting that the interactions between CrebA and Xbp1 may be more complicated and depend on the nature of the cell/organismal stressor. Further, ChIPseq data indicate that Creb3L regulation of Xbp1 transcription may be conserved in mammals (Consortium, 2012; Davis et al., 2018). Clearly, Creb3L family members play an important role in finely adjusting ER capacity to different environmental cues. However, there is more to be understood about their regulation and the links to the UPR.

To summarize, Creb3L-family transcription factors play key roles in regulating ER capacity in a variety of physiological contexts. Here, we specifically identify a clear and important function of this transcription factory family in mediating cellular and organismal responses to fasting and (re-) feeding in the liver, by directly impacting the expression and function of ER components. Our study indicates that this transcription factor family balances healthy responses to nutrients by regulating ER function and suppressing feeding. In metabolic disease, ER function is often severely disturbed (Ghemrawi et al., 2018). Maintaining the appropriate activity of these transcription factors may thus be important to establish a balance between “healthy” ER responses and “un-healthy” pathological responses, which contribute to metabolic diseases of the liver. Our study provides novel impetus for dissecting the vital links between transcriptional regulation, behavior and metabolic disease, thus impacting human health.

## ACKNOWLEDGEMENTS

We thank Andreas Ladurner, David Arnosti and Julia von Blume for scientific discussion and comments on the manuscript, Axel Imhof and Ignasi Forné for proteomics advice, as well as the Margulies team for discussion. We thank LMU LAFUGA for sequencing RNAseq and ChIPseq samples, the LMU BMC Proteomics Core for MS and Bioinformatics Core for training and advice on genomic and proteomic data analysis. We thank Teresa Burrell, Sonja Mühlberger and Fadila Benhamed for technical help and Kathrin Aschenbrenner for making fly food. We thank Deborah Andrew for CrebA antibody and fly lines, and Yan Li for the Fit mutant flies. We thank the Susan Eaton laboratory for Apoltp and Apolpp antibodies and Bruno Lemaitre for the Amy-p antibody. This research was supported by LMU, the Friedrich-Baur-Stiftung, the BMBF ERA-NET Neuron project *Food4Thought* and the Bavarian BioSysNet program. The EU Marie Skłodowska-Curie training network *ChroMe* (H2020-MSCA-ITN-2015-675610) funded H.K. and P.O.

## AUTHOR CONTRIBUTIONS

H.K. and C.M. designed and H.K. with the help of R.B. and M.T. performed the experiments. H.K. and C.M., analyzed and interpreted RNAseq, ChIPseq and MS results with the help from T.S.. Experiments with mice were designed by C.P. and performed by H.K. and P.O.. C.M. and H.K. wrote the manuscript with input from other authors. C.M. conceptualized and supervised the project.

## SUPPLEMENTAL FIGURE LEGENDS

**Figure S1:**
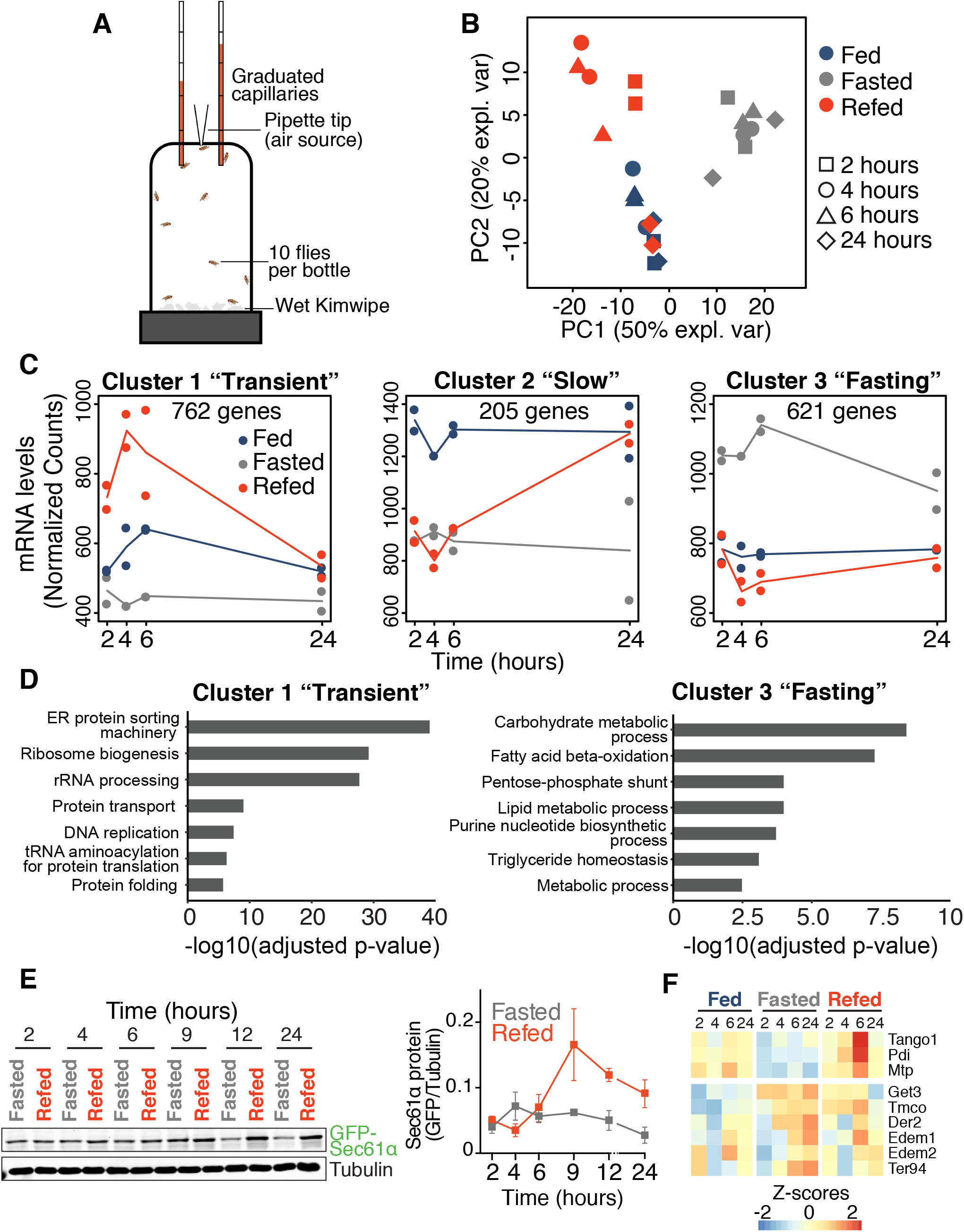
Feeding coregulates ER protein sorting machinery genes. **(S1A)** Schematic representation of the CAFÉ assay as previously described (Ja et al., 2007). **(S1B)** Principal component analysis showing the first two components for the mRNAseq data of the 1588 genes shown in (2E). **(S1C)** Using a second clustering approach using maSigPro identified 3 main clusters of the 1588 differentially expressed transcripts. **(S1D)** GO terms enrichment of cluster 1 and 3 using Fischer’s Exact test. **(S1E)** Representative western blot and quantitation of 3 independent western blots probing for Sec61α-GFP with a GFP antibody in refed and fasted flies. **(S1F)** A heatmap of mRNA levels of the machinery implicated in lipoprotein maturation and transport from the ER to the Golgi which change during refeeding compared to the mRNA other ER localized proteins which do not change in response to feeding.

**Figure S2:**
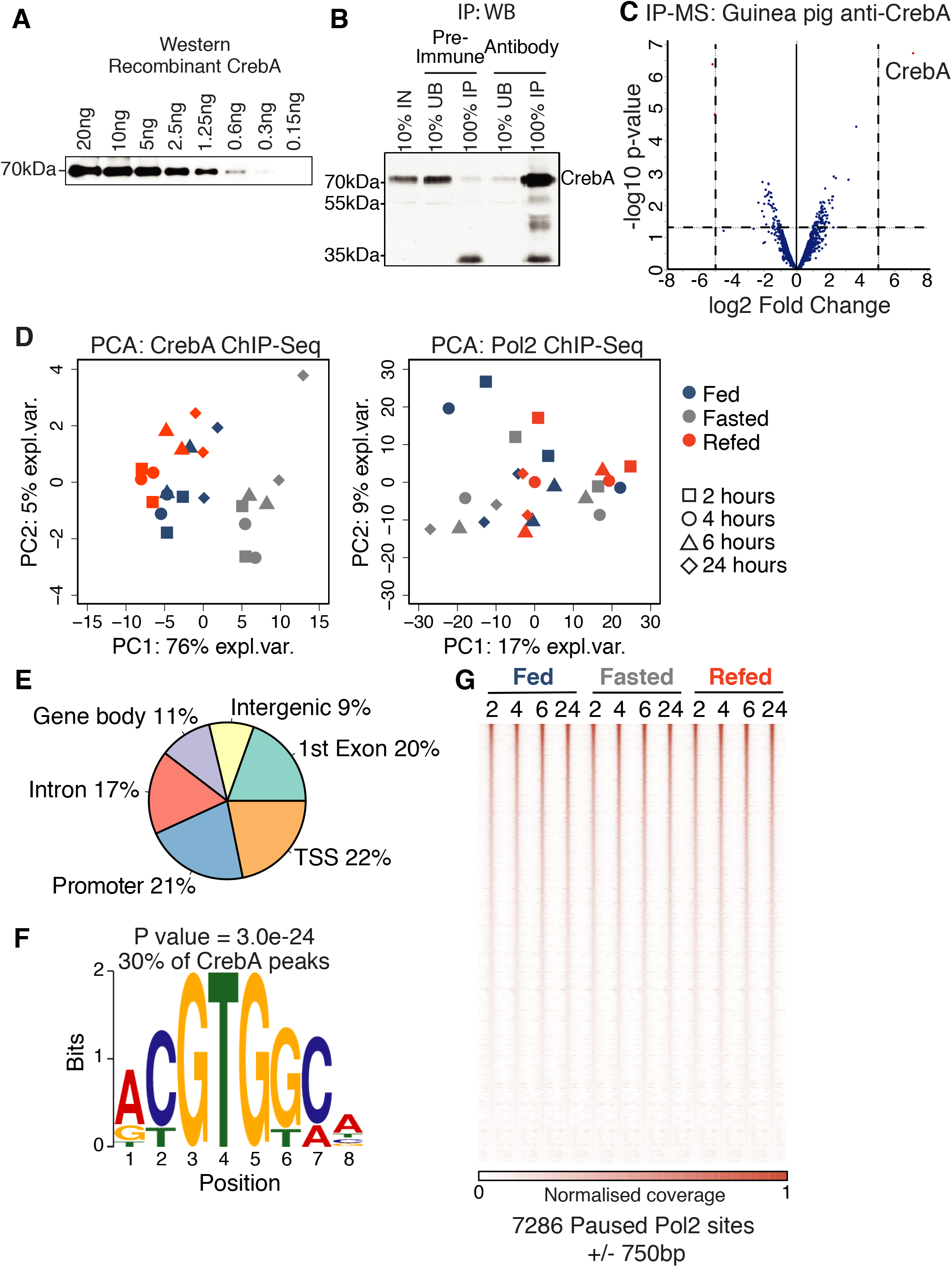
CrebA is a transcription factor regulated by feeding. **(S2A)** Representative western blot showing detection of recombinantly purified CrebA to sub nanogram levels using the anti-CrebA generated in Guinea Pig used for ChIP experiments. **(S2B)** Representative western blot showing immunoprecipitation (IP) of *Drosophila* embryo extracts using anti-CrebA antibody or with the preimmune serum as a control. **(S2C)** A volcano plot showing the quantification of IP experiments shown in (B) using LC-MS. Log Fold Change plotted representative for n=3 each pull downs either with the anti-CrebA antibody or preimmune serum. **(S2D)** Principal component analysis showing the first two components for CrebA ChIPseq (left) and Pol2 ChIPseq (right) data. Counts under respective peaks of each experiment were used to calculate the principal components. **(S3E)** Genomic distribution of 404 CrebA bound regions. **(S3F)** Enriched motif in the 404 CrebA binding sites. P-value was obtained using the MEME tool within MEMEChIP. 30% of total CrebA bound regions contain the motif. **(S3G)** A heatmap showing ChIPseq profiles of pull downs using an antibody against the Rpb3 subunit of Pol2 complex from female fly heads along the refeeding timecourse. Regions are +/−750bp of 7286 Pol2 called peaks were ordered in decreasing order based on the signal at refed 2-hours.

**Figure S3:**
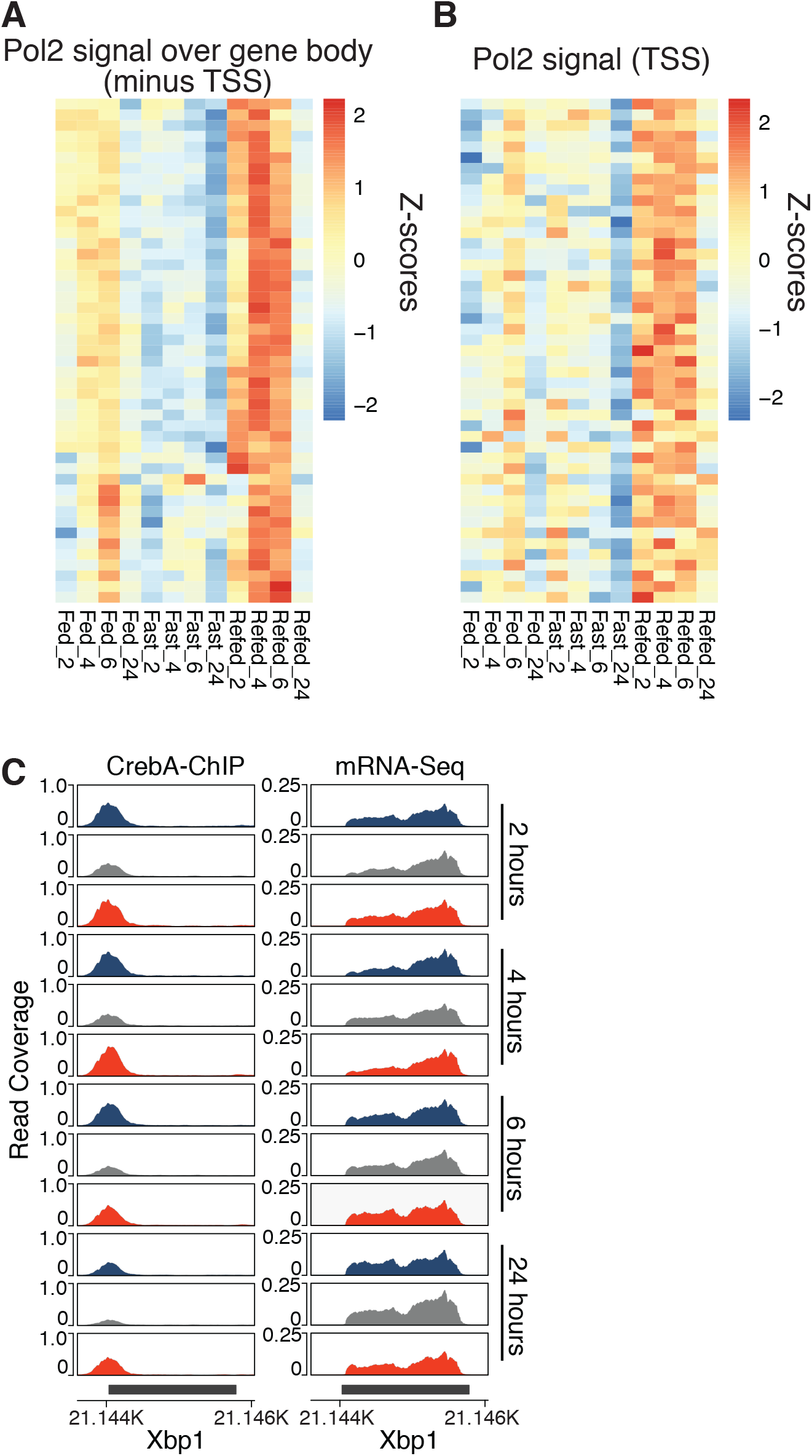
CrebA regulates the transcription of ER protein sorting machinery genes in response to feeding. Representative heatmap showing individual z-scores plotted for the 48 ER sorting machinery genes of Pol2 signal over gene body without the 200bp around the TSS **(S3A)** and only at the TSS **(S3B)**. Hierarchical clustering order same as in 5D. **(S3C)** Representative CrebA ChIPseq (left) and RNAseq (right) profiles from fly heads over Xbp1 – an example of a CrebA target gene that is not nutritionally responsive.

**Figure S4:**
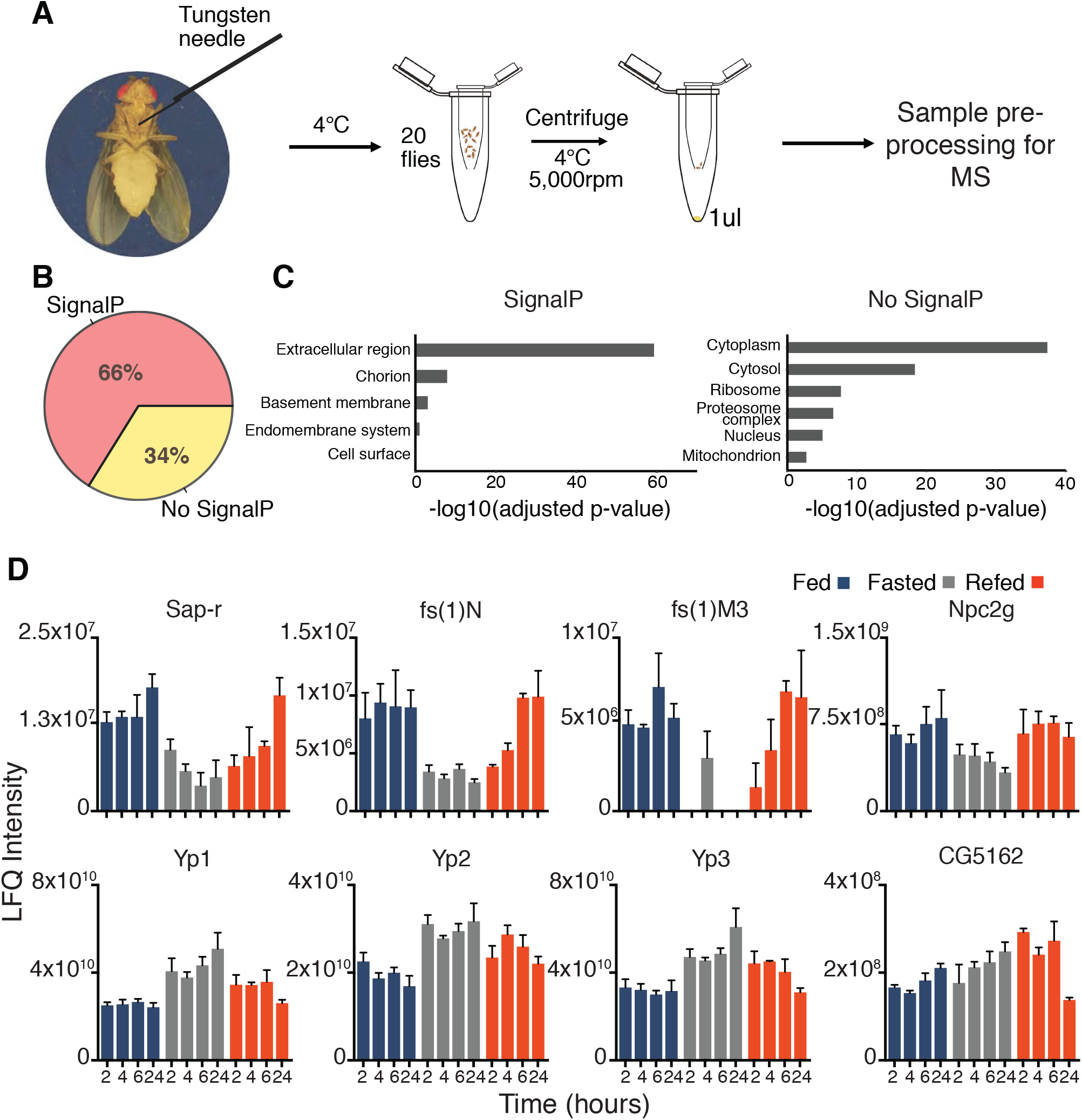
Hemolymph proteome changes with nutritional state. **(S4A)** An illustration showing the process of harvesting fly hemolymph. **(S4B)** Pie chart showing the percentages of the total 1878 proteins detected using MS predicted to have a signal peptide sequence. Prediction done using SignalP-5.0 (http://www.cbs.dtu.dk/services/SignalP/). **(S4C)** Functional enrichment of cellular component terms for proteins with or without a predicted signal peptide sequence using a Fischer’s Exact test. **(S4D)** Barplots showing levels of Sap-r, fs(1)N, fs(1)M3, Npc2g, Yp1, Yp2, Yp3 and CG5162 from the MS hemolymph data. LFQ normalized intensities plotted for n=3 ±SEM.

**Figure S5:**
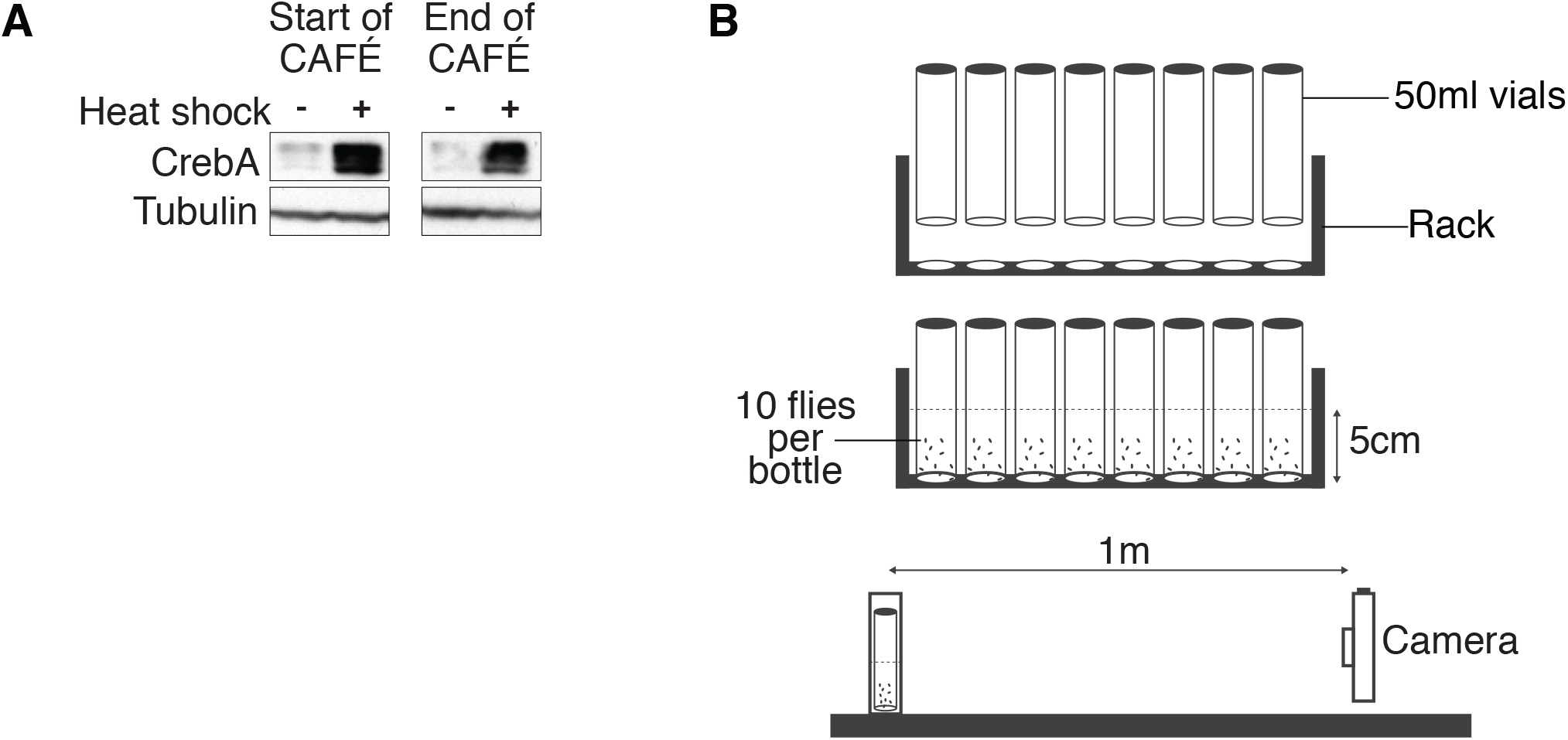
CrebA levels alter feeding and secreted lipoprotein in the hemolymph. **(S5A)** Representative western blot showing CrebA and H2A.Z protein levels in Hsp70-Gal4 crossed with UAS-CrebA flies right before and after measuring feeding. **(S5B)** Schematic diagram showing the working of a RING climbing assay.

**Figure S6:**
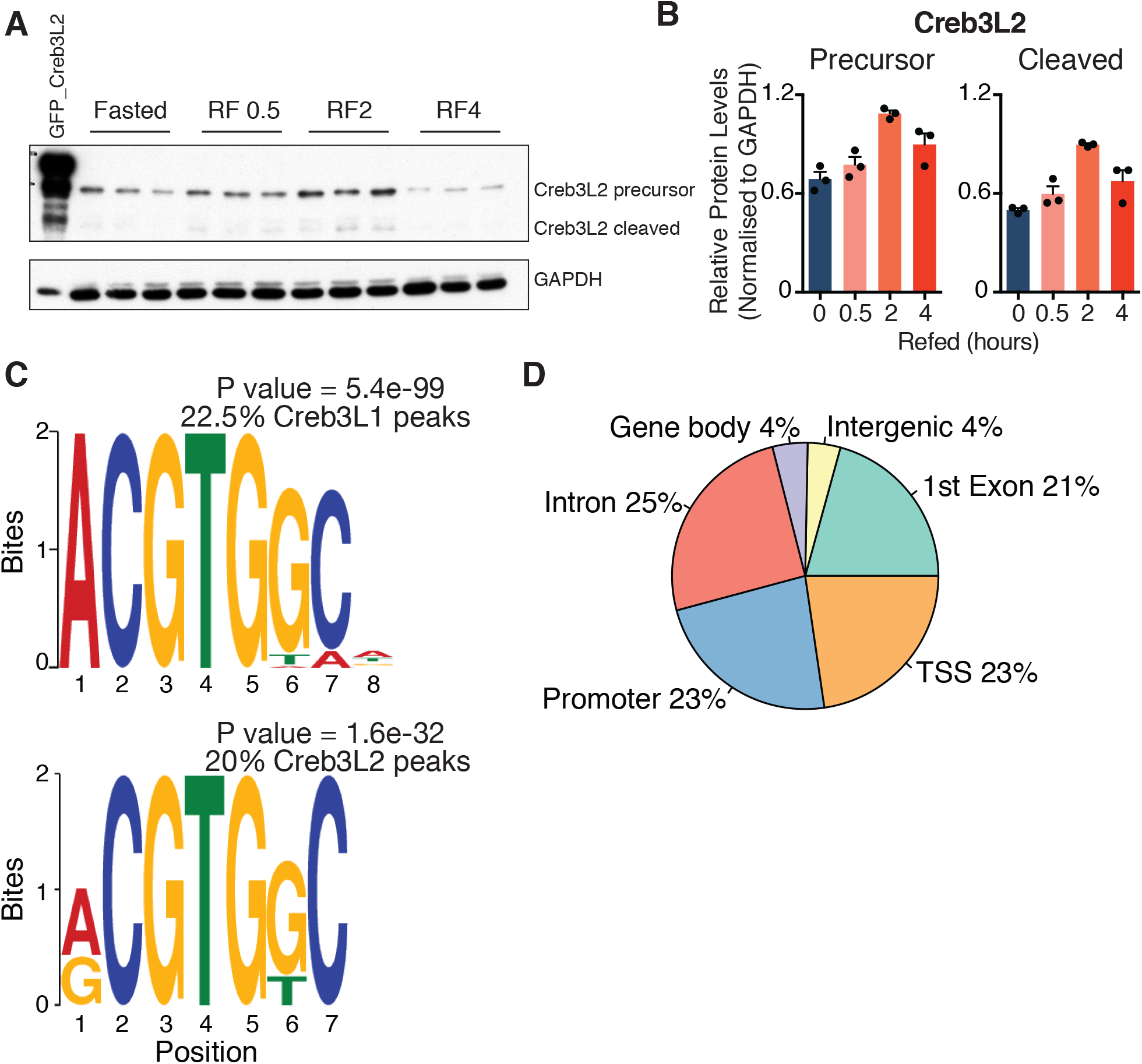
Mammalian homologs of CrebA regulate the early secretory machinery genes in response to feeding. **(S6A)** Western blot showing Creb3L2 protein levels of 3 additional biological replicates along the refeeding time-course in mice liver. **(S6B)** Quantification of the precursor and cleaved forms of Creb3L2 normalized to GAPDH shown in Figure 6D (n=3 ± SEM). **(S6C)** Motif found enriched in Creb3L1 or Creb3L2 bound regions using MEMEChIP. 20% of total peaks contain the motif using previously published ChIPseq data (Consortium, 2012; Davis et al., 2018) **(S6D)** Pie chart showing genomic distribution of all Creb3L2 bound regions in the ChIPseq data (Khetchoumian et al., 2019).

## SUPPLEMENTAL TABLES

**Table S1: Next-Generation mRNA Sequencing data, Related to Figure 1**

**Table S2: Peaks called for the ChIPseq experiments, Related to Figure 2**

**Table S3: Mass Spectrometry proteomic data, Related to Figure 4**

## STAR METHODS

### Lead contact

Further information and requests for resources and reagents should be directed to and will be fulfilled by the Lead Contact, Carla Margulies.

## EXPERIMENTAL MODEL AND SUBJECT DETAILS

### Fly strains

Flies were raised on standard sugar-yeast-agar medium. For the majority of experiments, the 2202U w^1118^, a Canton-S derivative strain was used (Boynton and Tully, 1992). Additional experiments were done using the Hsp70-Gal4 driver strain (Brand and Perrimon, 1993) and the UAS-CrebA strain (Rose et al., 1997). All analyses were performed on mated females. 3-5 days old flies were frozen in liquid nitrogen and stored at −80°C until used for further experiments. All experiments were done at 25°C, 60% humidity on a 12h light:12h dark cycle.

### Mouse models

C57BL/6N male mouse models obtained Jackson Laboratories were used for all experiments. Procedures were carried out according to the French guidelines for the care and use of experimental animals (animal authorization agreement number CEEA34.AFB/CP.082.12, Paris Descartes Ethical Committee). Mice were maintained in a 12-hr light: 12-hr dark cycle with free access to water and a standard chow diet (61% carbohydrate, 13.5% fat, and 25.2% protein; SAFE A03) unless otherwise specified.

### Cell lines

HEK293T cell lines (ATCC) were used. Cells were cultured at 37°C and 5% CO2 in high glucose DMEM media (Sigma) supplemented with 10% FCS (Gibco), 2mM glutamine (Gibco) and 1% penicillin/streptomycin (Gibco).

## METHOD DETAILS

### Capillary feeding assay (CAFÉ)

A modified capillary feeding assay (CAFÉ) was used to measure food intake and rate of consumption over time described previously (Ja et al., 2007). Briefly, 10 three to five days old flies were blown in to a single CAFÉ bottle with a hole on the side using an air pipette. A wet Kimwipe was placed on the bottom of the bottle as a source of water. Two graduated (5ul) capillaries (BLAUBRAND, Cat# BK8-708707) filled with 20% sucrose (Roth, Cat #4661.3), 5% yeast extract (Serva, Cat# 24540.3) and 5% (v/v) red dye (McCormick) were inserted at the top. Amount consumed was measured at desired time-points. Food consumption per fly (μL) was measured as food uptake divided by the total number of flies in a bottle at the end of the experiment. Rate of consumption (μL/hr) was calculated as food consumed in a single hour divided by the total number of flies in a single bottle.

### Fly climbing assay

A modified version of rapid iterative negative geotaxis (RING) assay described previously (Gargano et al., 2005) was used to measure fly climbing performance. Briefly, 10 flies were transferred to a 50mL vial marked with a 5cm height mark. 8 vials were together placed in a rack to hold the vials (Figure S5B). The rack together with the vials was tapped firmly to the bottom bringing all the flies to the bottom of the vials. Using a video camera, numbers of flies that climbed past the 5cm mark in 3 seconds were measured. Percentage climbed plotted was measured as the number of flies that climbed 5cm in 3 seconds over total number of flies in a vial.

### RNA extraction

Thirty female fly heads were homogenized in Trizol reagent (Life Technologies, Cat#15596018) and left at RT for 5 minutes. The lysates were extracted once with chloroform and spun down at 12000g, 4C for 10 minutes. The upper aqueous phase was taken and RNA was precipitated with isopropanol and 1uL of 200mg/ml glycogen (Thermofischer, Cat#10814010). Pellets were washed twice with 75% ethanol, air-dried and resuspended in 30ul RNAse free dH20.

### cDNA preparation

15ug of RNA was taken and treated with 0.5uL TURBO DNA-free DNAse 1 (Ambion, M1907) at 37°C for 30 minutes. The reaction was inactivated using 6uL of DNAse Inactivation Reagent as described in manufacturer’s instructions. For cDNA synthesis, 1ug of DNAse-free RNA was reverse transcribed using random primers (Thermo, 48190011) and 1uL SuperscriptIII (Thermo, 18080085) according to manufacturer’s instructions.

### RTqPCR

cDNAs were quantified by qPCR using FAST or POWRUP SYBR Green Master Mix (Life Technologies) and a 7500 Fast or QauntStudio™ 3 Real-Time PCR systems. cDNA was amplified using gene specific primers (Table) and normalized based on a standard curve generated against genomic DNA extracted from wilt-type flies. Absolute values were plotted normalized to H2AZ.

### Transcriptomics

RNA was extracted as described above. A sequencing library was prepared using Illumina polyA-mRNA library preparation methods with paired-end option. Both library preparation and sequencing were performed by the EMBL Genomics Core Facility. Fifty base pair reads were obtained from an Illumina HiSeq 2000 sequencer. Reads were aligned to the reference genome (version dm6) using STAR (version 2.6.0). Uniquely mapped reads were counted per genes using STAR-quantMode GeneCounts using the annotation dm6.13. Differential expression analysis was performed using maSigPro (version 3.6). Coverage vectors were calculated from BAM files (generated by STAR) using tstools (https://github.com/musikutiv/tsTools/) and normalized over total coverage. Further analysis was performed using R and graphs were plotted using R base graphics.

### Informatic processing of the transcriptomics

Reads were aligned to the reference genome (version dm6) using STAR (version 2.6.0). Uniquely mapped reads were counted per genes using STAR-quantMode GeneCounts using the annotation dm6.13. Coverage vectors were calculated from BAM files (generated by STAR) using tstools (https://github.com/musikutiv/tsTools/) and normalized over total coverage. Read counts obtained from STAR were normalized using trimmed means of M values using the NOISeq package (version 2.2) in R. Differential expression analysis was performed using maSigPro (version 3.6). A significance level cutoff of 0.05 and a R-squared value cutoff of 0.6 was used for variable selection in the stepwise regression models. Hierarchal clustering based on Euclidean distance was performed within maSigPro to identify 3 clusters in Figure S1C. Significant genes called were used to plot the heatmap in Figure 1E. GO term analysis was done using a Fischer’s Exact Test and the database from biomaRt (version 2.38) in R.

### Hemolymph extraction

Hemolymph was extracted as previously described (http://musselmanlab.com/wp-content/uploads/2018/09/adult-hemolymph-isolation-and-sugar-assays.pdf). Briefly, 20 flies immobilized on ice were pricked in the thorax with a (diameter) tungsten needle (Musselman et al., 2013). A 0.5mL eppendorf tubes with three 0.25um holes at the bottom was placed in a bigger 1.5mL eppendorf tube. Flies were transferred to the smaller Eppendorf and the whole apparatus was centrifuged for 5min, 5000rpm at 4°C to collect 1uL of hemolymph.

### Mass spectrometry

#### Peptide preparation

Peptides from hemolymph samples were prepared using the Preomics iST sample preparation kit (Cat# P.O.00027) as per manufacturer’s instructions. For IP experiments samples were trypsin digested and peptides were desalted using C°18 Stagetips.

#### LC-MS/MS

Samples were evaporated to dryness, resuspended in 15 μl of 0.1% formic acid solution and injected in an Ultimate 3000 RSLCnano system (Thermo), either separated in a 25-cm Aurora column (Ionopticks) with a 50-min gradient from 6 to 43% of 80% acetonitrile in 0.1% formic acid (hemolymph samples) or separated in a 15-cm analytical column (75μm ID with ReproSil-Pur C18-AQ 2.4 μm from Dr. Maisch) with a 50-min gradient from 5 to 60% acetonitrile in 0.1% formic acid (IP samples). The effluent from the HPLC was directly electrosprayed into a Qexactive HF (Thermo) operated in data dependent mode to automatically switch between full scan MS and MS/MS acquisition. Survey full scan MS spectra (from m/z 375–1600) were acquired with resolution R=60,000 at m/z 400 (AGC target of 3×10^6^). The 10 most intense peptide ions with charge states between 2 and 5 were sequentially isolated to a target value of 1×10^5^, and fragmented at 27% normalized collision energy. Typical mass spectrometric conditions were: spray voltage, 1.5 kV; no sheath and auxiliary gas flow; heated capillary temperature, 250°C; ion selection threshold, 33.000 counts.

### LC-MS/MS data analysis

#### MaxQuant

Raw MS data files were processed using MaxQuant (version 1.6.3.4) with the Andromeda search engine with FDR < 0.01 at protein and peptide level. The default settings were used with the following modifications: variable modification methionine (M), acetylation (protein N-term) and the fixed modification carbamidomethyl (C) were selected, only peptides with a minimal length of seven amino acids were considered. Peptide identification was done using the drosophila melanogaster DB from Uniprot (uniprot_3AUP000000803_Drosophila_melanogaster_20180723.fasta).

#### Perseus

Further analysis was done using Perseus (version). Contaminants and reverse hits were removed. Label-Free Quantification (LFQ) normalized intensity values were taken for proteins containing equal to or greater than 8 out of 12 non-zero values for at least 2 out of the 3 conditions (Fed, Fasted and Refed). Missing values were imputed based on a Gaussian distribution using default parameters.

#### R

Imputed and normalized LFQ values were imported in R. Fasta sequences were downloaded for all detected proteins from Uniprot and signal peptide prediction was done using SignalP 4.0. PCA analysis was done and visualized using R base functions and graphics. Differential expression analysis was performed using maSigPro.

### Generation of anti-CrebA antibody

Guinea-pig polyclonal anti-CrebA antibodies were generated against the whole CrebA protein (clone from Drosophila Genomics Resource Center, 1623052). The recombinant protein was expressed in a pETM11 vector in E.Coli and purified using a histidine tag column. Histidine tag was removed using a TEV cleavage site in between the CrebA protein and the histidine tag. Purified antigen was shipped to Eurogentec, Belgium where the animals were immunized and sera was collected under contract number DE17051.

### Western Blotting

Proteins were separated using 10% SDS-PAGE gels at constant voltage of 200 V. Transfer of proteins to a nitro-cellulose membrane was carried out at constant 100 V after which the membranes were blocked in 5% milk powder prepared in TBS-T solution for 1 hour. Primary antibody incubation was done over-night and membrane washed 3 times for 10 minutes in TBS-T. Secondary antibody incubation was done for 1 hour after which the membrane was washed again for 3 times for 10 minutes. Proteins were visualized by adding a 1:1 HRP substrate solution (Cat) with a developer. All primary and secondary antibodies were diluted with 5% milk containing TBS-T. 5 fly heads were loaded for all westerns shown. Other than the Guinea-Pig anti-CrebA antibody generated, rabbit polyclonal anti-CrebA (1:200, DSHB) and anti-H2AZ (1,5:000) (Schauer et al., 2013) were used.

### Immunoprecipitation

Immunoprecipitation assays were performed using Drosophila embryo extracts prepared as previously described (Becker and Wu, 1992). Briefly, 100 grams of embryos were collected, washed and homogenized in homogenization buffer [15mM Hepes pH 7.6, 10mM KCl, 5mM MgCl2, 0.5mM EGTA pH 8.0, 0.1mM EDTA pH8.0 supplemented fresh with 1mM DTT, 0.2mM PMSF, 1mM NaMBS, 1ug aprotinin, 1ug leupeptin, 1ug pepstatin]. Nuclei were pelleted at 10,000g for 15 minutes and components of the nuclei precipitated using (NH_4_)_2_SO_4_. 1ul of antibody was incubated overnight with the 500ug of the extract. 20ul of washed protein A Dynabeads were added and incubated at 4°C for 3 hours. Beads were washed on a magnetic rack. Fraction of the reaction was used for western blots while the rest was further processed for mass spectrometry. Pre-immune sera from the same animals was used as a control.

### ChIPseq

ChIP experiments were done as previously described in Tamas et al., 2013. Briefly, 1000-1500 heads were separated from frozen flies using 630 and 400 microns sieves and homogenized in homogenization buffer [350 mM sucrose, 15 mM HEPES pH 7.6, 10 mM KCl, 5 mM MgCl2, 0.5 mM EGTA, 0.1 mM EDTA, 0.1% Tween, with 1 mM DTT and Protease Inhibitor Cocktail (PIC) (Roche) added immediately prior to use] at 4°C. Solution was fixed using 1% formaldehyde and quenched with glycine. Nuclei were filtered using a 60 microns nylon filter, washed 3 times with RIPA (150 mM NaCl, 25 mM Tris-HCl pH 7.6, 1 mM EDTA, 1% Triton-X, 0.1% SDS, 0.1% DOC, with protease inhibitors added prior to use) and sonicated using Branson 250 (2 cycles, intensity 5, pulsing 60 s) and Covaris S220 (PIP150, DC20, CPB200, time 10 min) sonicator. Soluble chromatin was taken as supernatant after centrifugation and stored at −80°C.

Ten ug of chromatin was used for a single ChIP experiment. Chromatin was pre-absorbed with Sepharose protein A beads equilibrated with RIPA buffer containing 1 μg/μl salmon sperm DNA and 1 μg/μl BSA. Chromatin was incubated with the relevant antibody overnight. 10% input material was separated at this point and later pooled to form one sample per condition/time-point. Pull-downs were performed using the equilibrated beads. Antibodies used were anti-CrebA (Guinea-Pig, own stock), anti-RPB3 (Schauer et al., 2013). Immunoprecipitated DNA was purified using AGENCOURT AMPURE XP magnetic beads (Beckmann-Coulter). At least 2 biological replicates were used for each experiment. In some cases, additional technical replicates were done in order to obtain the required amount of chromatin.

The ChIPseq libraries were prepared with 1 ng of ChIP and input DNA with NEBNext®Ultra II DNA Library Prep Kit for Illumina® according to the manual instructions. The libraries were barcoded using NEBNext® Multiplex Oligos Set 1 and 2 and sequenced at LAFUGA at the Gene Center (LMU) using an Illumina® HiSeq1500 sequencer.

### ChIPseq analysis

ChIPSeq 50bp single end reads were aligned to the reference *Drosophila melanogaster* (dm6) using bowtie2 (version 2.2.9). Peak calling was done using macs2 (version 2.1.1) with a fold change cut-off of 10. Input was used as a control. Reads under the regions were calculated using subread (version 1.6.2). Regions were associated to nearest genes using RGmatch. For the peak regions heatmap analysis, coverages for +/− 750bp from peak summit were extracted and plotted using R base functions. GO term analysis was done using a Fischer’s Exact Test and the database from biomaRt (version 2.38) in R. Bound CrebA was calculated as number of reads present in peaks closest to genes. Pol2 over gene body was calculated as the number of reads over the entire gene minus the TSS (first 200bp).

### Mice experiments

#### RNA and ChIPseq analysis

Count matrix was downloaded from the submitted repository GSE118973 and analyzed using maSigPro. Creb3L1 ChIPseq was downloaded from encodeproject.org with the accession ENCFF950GGY. Peaks from the BAM files were called using macs2 (version 2.1.1) with a fold change cut-off of 7. Regions were associated to nearest genes using RGmatch. GO term analysis was done using a Fischer’s Exact Test with the GO terms database from biomaRt (version 2.38) in R.

#### Mice liver analysis

Mice were fasted for 24 hours and refed with standard chow diet plus 20% glucose solution for 0, 0.5, 2 or 4 hours. The sacrifice was always performed at noon in all groups to avoid circadian effects. Livers collected were snap-frozen and stored at −80°C. RNA from 15mg of mice liver was extracted and cDNA prepared as described above. RTqPCR was done using primers amplifying small regions of Creb3L1 and Creb3L2. Data was normalized to the house-keeping gene TATA-binding protein (TBP). For westerns, whole-cell protein lysates were prepared from 20mg of liver and analyzed as described above. Mouse monoclonal anti-Creb3L2 (Merck, MABE1018) and anti-GAPDH (Genetex, GT239) antibodies were used.

#### Cell culture experiments

Creb3L2 coding sequence was cloned out from mice liver cDNA and introduced in a pEGFP-C1 (Clonetech) vector. HEK293T cells were transfected with 10ug of vector and protein lysates were prepared 2 days after. GFP tagged Creb3L2 expression was confirmed under a microscope and western blot analysis using antibodies against GFP (developed in-house).

## QUANTIFICATION AND STATISTICAL ANALYSIS

Statistical analyses were carried out using GraphPad Prism 9 and methods are descripted in the legend of the figures or in the methods section for the respective technique.

## DATA AND CODE AVAILABILITY

RNA and ChIP sequencing data are deposited in the National Center for Biotechnology Information Gene Expression Onmibus (https://www.ncbi.nlm.nih.gov/geo/), accession number GEO: GSE165547. The mass spectrometry proteomics data have been deposited to the ProteomeXchange Consortium via the PRIDE (Perez-Riverol et al., 2019) partner repository with the dataset identifier PXD023886.

